# Genetic diversity, virulence, antimicrobial resistance genes, and antimicrobial susceptibility of group B *Streptococcus* (GBS) linked to mass mortalities of cultured Nile tilapia in Brazil

**DOI:** 10.1101/2022.12.25.521911

**Authors:** Inácio Mateus Assane, Rubens Ricardo de Oliveira Neto, Daniel de Abreu Reis Ferreira, André do Vale Oliveira, Diogo Teruo Hashimoto, Fabiana Pilarski

## Abstract

*Streptococcus agalactiae*, group B *Streptococcus* (GBS), is the main bacterial pathogen of cultured Nile tilapia in many countries and causes extensive deaths in all stages of the farming cycle throughout the year. This study investigated the genetic diversity, virulence, presence of antibiotic resistance genes and antimicrobial susceptibility of 72 GBS linked to mass mortalities of cultured Nile tilapia in Brazil. Isolate identity was confirmed by morphological, biochemical and molecular analyses. Capsular serotype, multi-locus sequence typing (MLST) allelic profiles and putative pathogenic factors were identified by polymerase chain reaction (PCR), gel electrophoresis, sequencing and molecular analyses of different genes. The presence of antimicrobial resistance genes and antimicrobial susceptibility to florfenicol (FFC), oxytetracycline (OTC), thiamphenicol (TAP) and their combination were evaluated by PCR, followed by gel electrophoresis, and broth microdilution antimicrobial susceptibility testing, respectively. All clinical isolates studied were confirmed to be GBS, one from serotype III (IA2022) and 71 from serotype Ib, suggesting that serotype Ib was the most prevalent strain between 2011 and 2016 in the south and southern and southeastern regions of Brazil. Eight different allelic profiles were identified for the first time, being *adhP-* 52, *pheS-*2, *atr-*31, *glnA-*4, *sdhA-*2, *tkt-*19 the most predominant. Between one (*glcK*) and three (*adhP* and *glnA*) alleles were present at each locus. All strains, except IA2022, showed a partial gene deletion event on the *glcK* gene. The surface protein *Rib* and hypervirulent GBS adhesin *BibA* were detected in all strains, except for 18P, which was negative for *rib*. On the other hand, α and β antigens of the C protein were only detected in IA2022. All antimicrobials showed high minimum inhibitory concentration (MIC ≥ 16 µg/mL) values against several strains with negative results for resistance genes. Despite indifference and antagonism being the most predominant activities in all combinations evaluated, the record of synergism, including in a strain with a resistance gene and phenotypic resistance, suggests that combination therapy can have therapeutic efficacy when well planned. The combination involving OTC and TAP or FFC is a likely candidate for improving the treatment of streptococcosis using combination therapy, even for strains showing phenotypic and genotypic resistance to OTC. This study provides pertinent data on pathogenic GBS genetic diversity, virulence, the presence of antibiotic resistance genes and antimicrobial susceptibility, which may be useful in the development of effective vaccines and therapeutic strategies for the prevention and control of streptococcosis in aquaculture farms.

## 1. Introduction

Nile tilapia, *Oreochromis niloticus*, is one of the major species produced in the world aquaculture. According to the Food and Agriculture Organization of the United Nations (FAO, 2022), in 2020, it was the third major species produced, after grass and silver carps (*Ctenopharyngodon idellus* and *Hypophthalmichthys molitrix*, respectively), sharing 9% of finfish production. Due to its zootechnical characteristics, such as strong environmental adaptability, ease of breeding, disease resistance, high protein content, large size, rapid growth and palatability, Nile tilapia is probably the most widely introduced species and economically important food fish in many countries, being introduced in 114 countries or regions (FAO, 2021), including Brazil, where it represented, 62.3% of fish production in 2020 (IBGE, 2020).

Brazil has a good tilapia sector, which grows fast and represents over 74% of the production of Nile tilapia in the Americas. In the last decade, tilapia production increased from 150,000 to over 400,000 tonnes, making Brazil the fourth major Nile tilapia producer in the world, behind China, Indonesia and Egypt (FAO, 2021; IBGE, 2020; Peixe BR, 2021). However, this sector is being negatively impacted by bacterial diseases (Assane, Prada Mejia, et al., 2021; Chideroli et al., 2017; Delphino et al., 2019; Junior et al., 2020; Sebastião, Furlan, Hashimoto, & Pilarski, 2015; Sebastião, Pilarski, Kearney, & Soto, 2017) and the lack of effective therapies (Assane, de Sousa, et al., 2021; de Oliveira, Queiroz, Teixeira, Figueiredo, & Leal, 2018). According to Tavares-Dias and Martins (2017), in Brazil, the annual loss to the inland aquaculture sector due to parasitic and bacterial diseases is estimated to be USD 84 million.

Currently, *Streptococcus agalactiae*, group B *Streptococcus* (GBS), is the main bacterial pathogen of Nile tilapia and causes extensive deaths in all stages of the farming cycle throughout the year (Chideroli et al., 2017), with higher frequency when the water temperature is above 28 °C (Delphino et al., 2019). On the other hand, only two antimicrobials (florfenicol-FFC and oxytetracycline-OTC) are permitted for use in Brazilian aquaculture (SINDAN, 2022), despite the reports of vaccine failure and antibiotic ineffectiveness after a streptococcosis outbreak (Chideroli et al., 2017; de Oliveira et al., 2018).

Isolates of GBS show distinct genetic diversity, as well as variability in biochemical and physiological characteristics. Knowledge about these characteristics is necessary for the correct diagnosis, development and implementation of effective prevention and treatment strategies. The polysaccharide capsule (CPS), encoded by the *cps* gene (Imperi et al., 2010); the allelic profile of seven different loci (sequence type (ST)) (Jones et al., 2003), which is used to group strains into clonal complexes (CC) (Honsa et al., 2008); and the surface proteins (virulence proteins), such as the α and β antigens of the C protein (*bca* and *bac*, respectively), *BibA*, C5a peptidase (*scpB*), hyaluronate lyase (*hly*), *hvgA* and *Rib* (Rajagopal, 2009) are the main sources of its diversity. There are ten structurally and serologically distinct CPS types (capsular serotypes: Ia, Ib and II-IX), more than 1800 ST (Jolley, Bray, & Maiden, 2018), several clonal complexes (CC) (Honsa et al., 2008) and putative virulence factors that have been identified and associated with a strain’s virulence (Marques, Kasper, Pangburn, & Wessels, 1992; Rajagopal, 2009) and host’s immune response (Bianchi-Jassir et al., 2020; Flaherty et al., 2019). Thus, capsular serotype, ST, CC and virulence factors identification are necessary for both epidemiological studies and the development of effective prevention and treatment strategies.

To date, five capsular serotypes (Ia, Ib, II, III and IV) and six CC (CC7, CC19, CC23, CC103, CC283, CC552) have been linked to disease outbreaks in cultured fish in different countries (Chen et al., 2015; Delannoy, Samai, & Labrie, 2021; Sudpraseart, Wang, & Chen, 2021), including Brazil (Chideroli et al., 2017; Delphino et al., 2019; Godoy et al., 2013). Current data indicate that in Brazil, capsular serotype Ib and III, and ST-103, ST-260, ST-552 and ST-553 (CC552) are the most prevalent, although strains belonging to capsular serotype Ia and non-serotypeable strains have been isolated from disease outbreaks (Barony et al., 2017; Chideroli et al., 2017; Godoy et al., 2013). However, information about the geographical distribution of sequence types, virulence, antimicrobial susceptibility profile and the mechanisms of antibiotic resistance of GBS isolated from Nile tilapia farmed in Brazil is still scarce.

This study aimed to enrich our current knowledge in GBS epidemiology and provide pertinent data for the development of effective GBS infection prevention and treatment methods. Here, we present the genetic diversity, virulence, mechanisms of antibiotic resistance and antimicrobial susceptibility of GBS linked to mass mortalities of cultured Nile tilapia in Brazil. Capsular serotype, ST and putative pathogenic factors were identified. The presence of antimicrobial resistance genes and antimicrobial susceptibility to FFC, thiamphenicol and their combination were evaluated.

## 2. Materials and methods

### 2.1. Clinical isolates

Seventy-two clinical isolates of *Streptococcus agalactiae* (group B Streptococcus, GBS) were recovered between 2011 and 2018, from the brain, liver and kidney of diseased Nile tilapia, *Oreochromis niloticus*, that were farmed in different locations in Brazil (Figure 1) and used to determine the genetic diversity, virulence, the presence of antimicrobial resistance genes and antimicrobial susceptibility to florfenicol (FFC), oxytetracycline (OTC), thiamphenicol (TAP) and their combinations.

**Figure 1.**
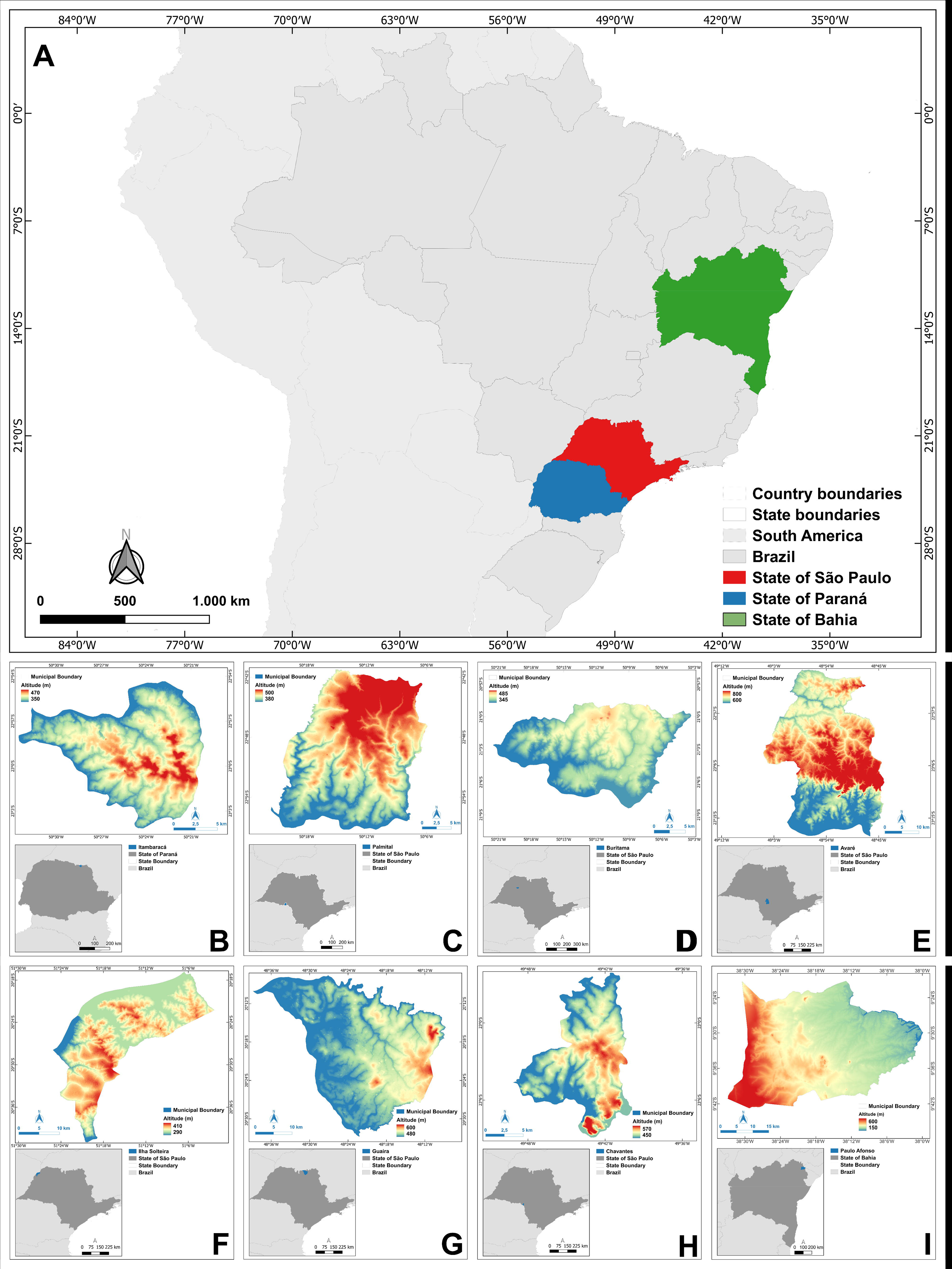
Map showing the geographical location of clinical isolates of *Streptococcus agalactiae* (group B *Streptococcus*, GBS) recovered between 2011 and 2018, from the brain, liver and kidney of diseased Nile tilapia, *Oreochromis niloticus*, farmed in Brazil. Datum SIRGAS 2000. Data source: IBGE, 2021; SRTM/NASA (Earth Explorer USGS).

### 2.2. Genetic diversity

The genetic diversity was determined by a polymerase chain reaction (PCR) and the sequencing of different genes used for identification, multi-locus sequence typing (MLST), capsular serotyping, and detection of virulence and antibiotic resistance factors in GBS. As described in Table 1, several genes were amplified, such as seven housekeeping genes, alcohol dehydrogenase (*adhP*), phenylalanyl tRNA synthetase (*pheS*), amino acid transporter (*atr*), glutamine synthetase (*glnA*), serine dehydratase (*sdhA*), glucose kinase (*glcK*), and transketolase (*tkt*); the virulence genes, capsular polysaccharide (*cps*), β antigens of the C protein (*bac*), α antigens of C protein (*bca*), hyaluronate lyase (*hly*), surface protein Rib (*rib*), C5a peptidase (*scpB*), and hypervirulent GBS adhesins (*hvgA* and *BibA*); and the antibiotic resistance genes, the mobile genetic element (MGE) insertion sequence (IS) 1548; erythromycin Ribosomal Methylase (*ermB* and *ermTR*), erythromycin resistance efflux pump (*mefA*), lincomycin and clindamycin resistance gene-lincosamide nucleotidyltransferase (*linB*), tetracycline resistance proteins (*tetM* and *tetO*), and phenicol exporters (*fexA* and *fexB*) were amplified as described in Table 1. MLST allelic profiles and capsular serotype of each clinical isolate were determined as previously described by Jones et al. (2003) and Imperi et al. (2010), respectively.

**Table 1.**
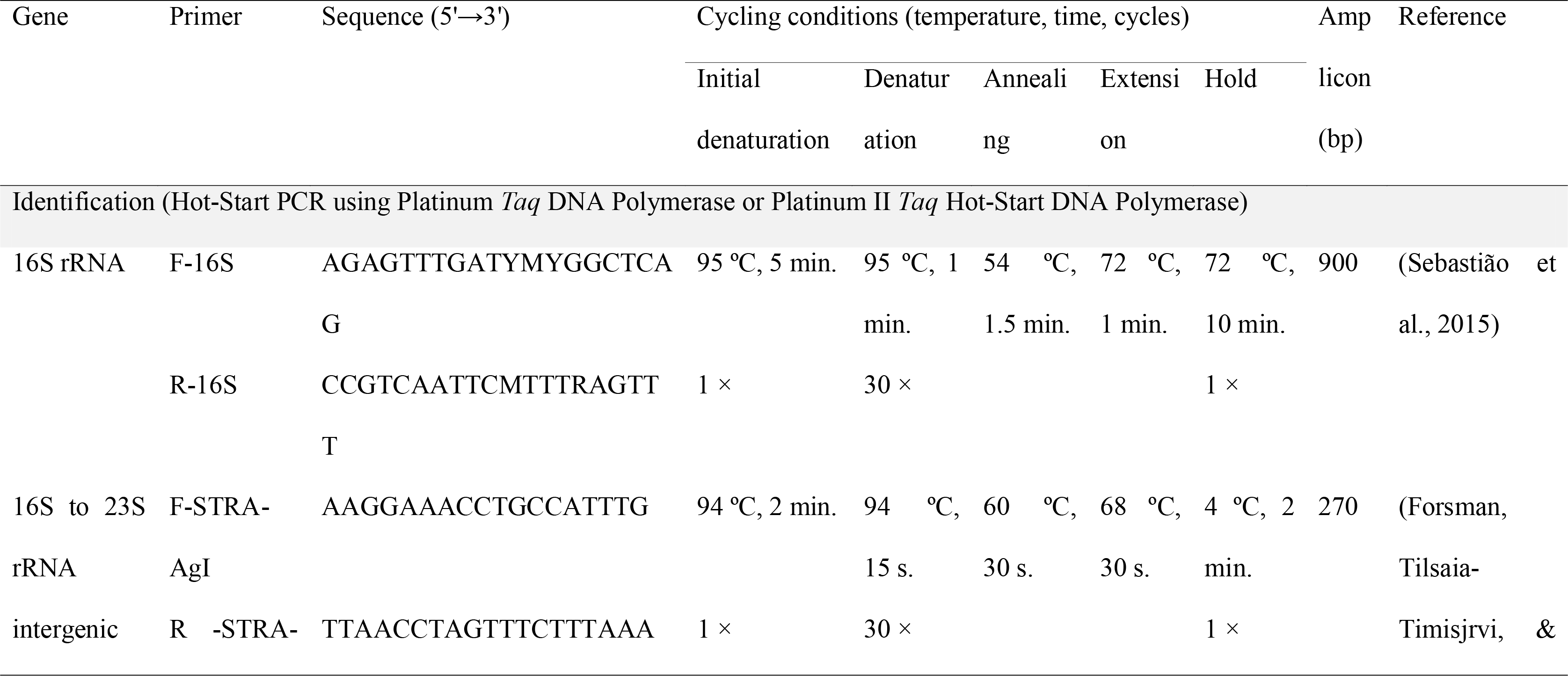
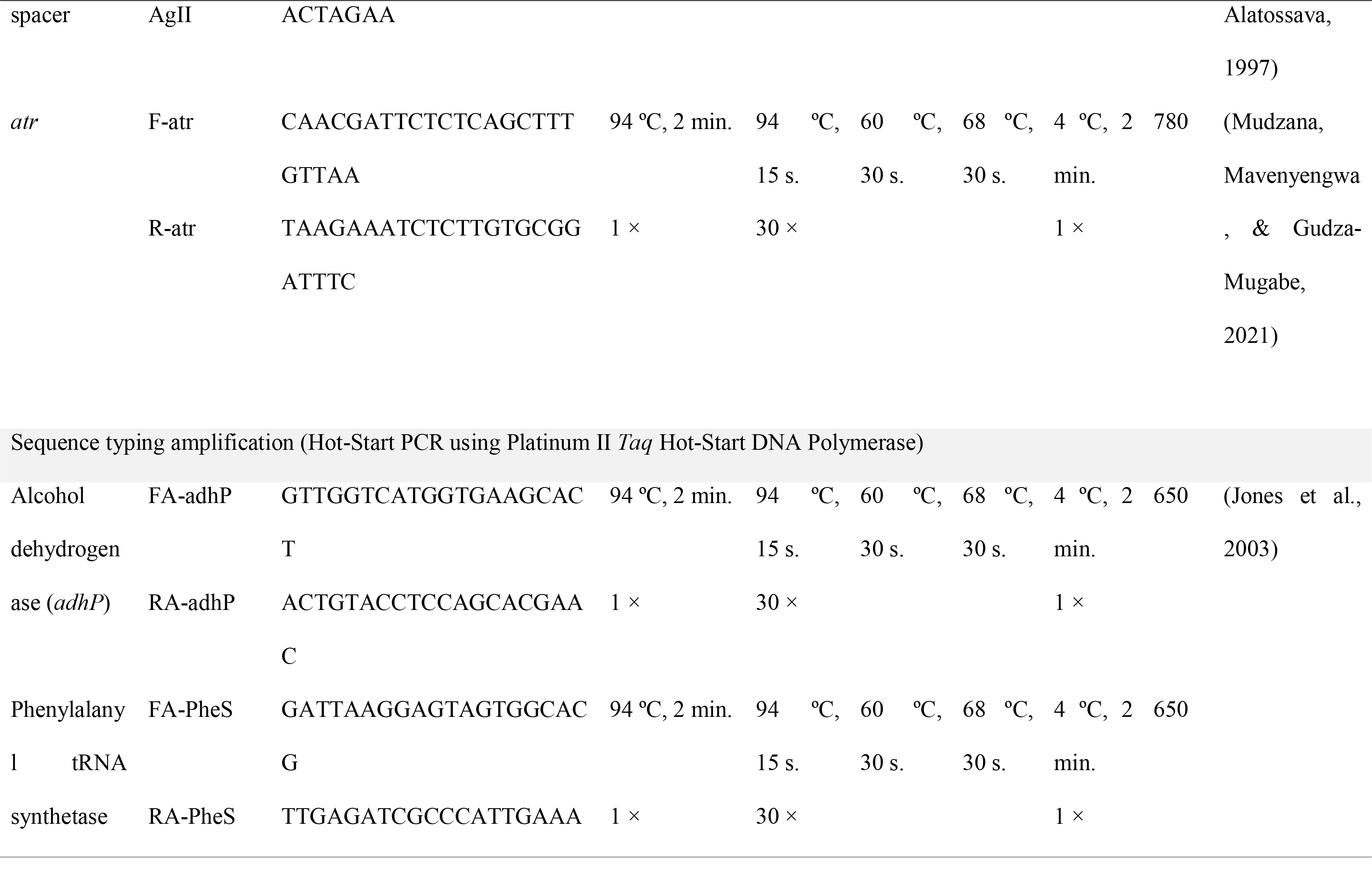
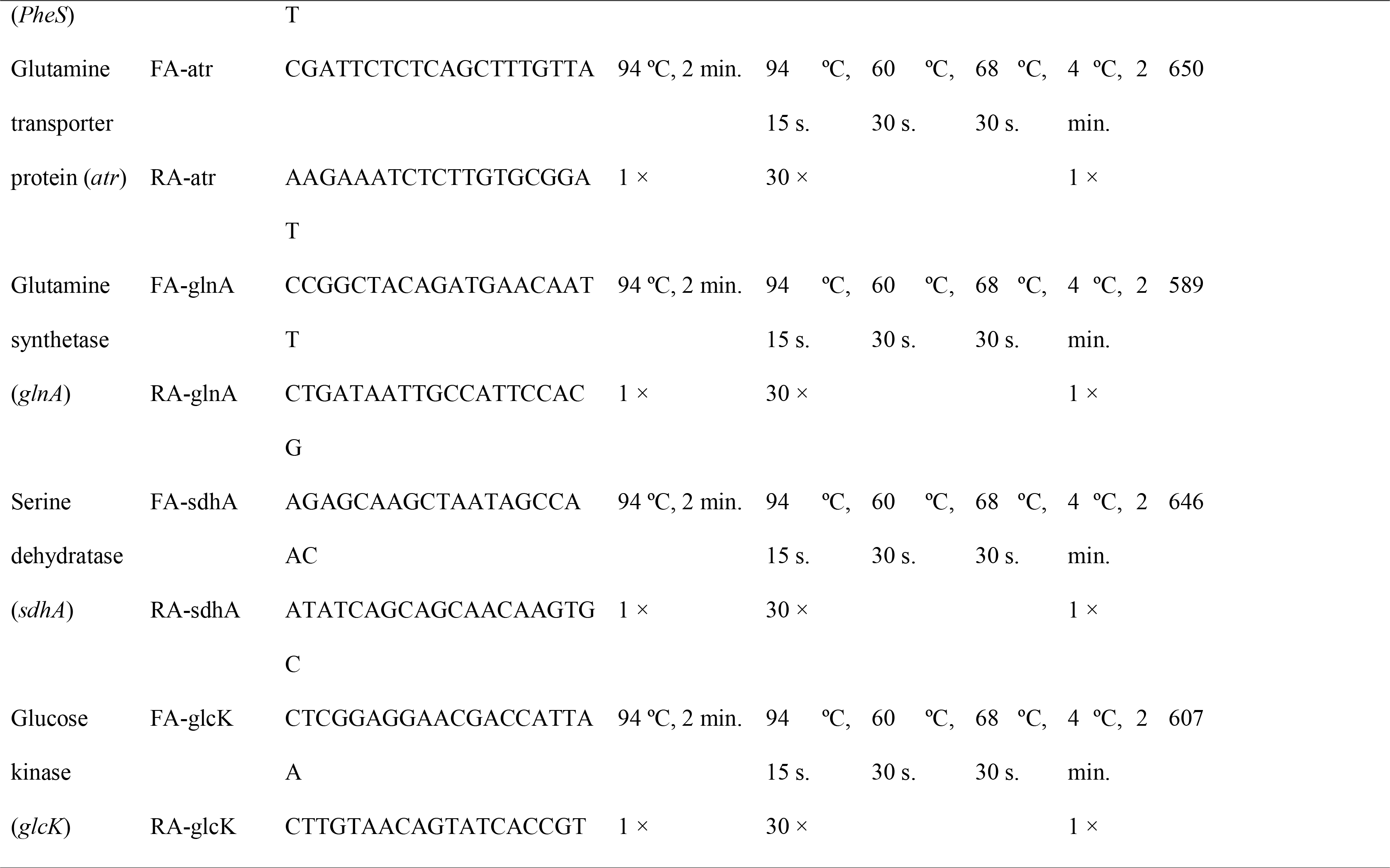
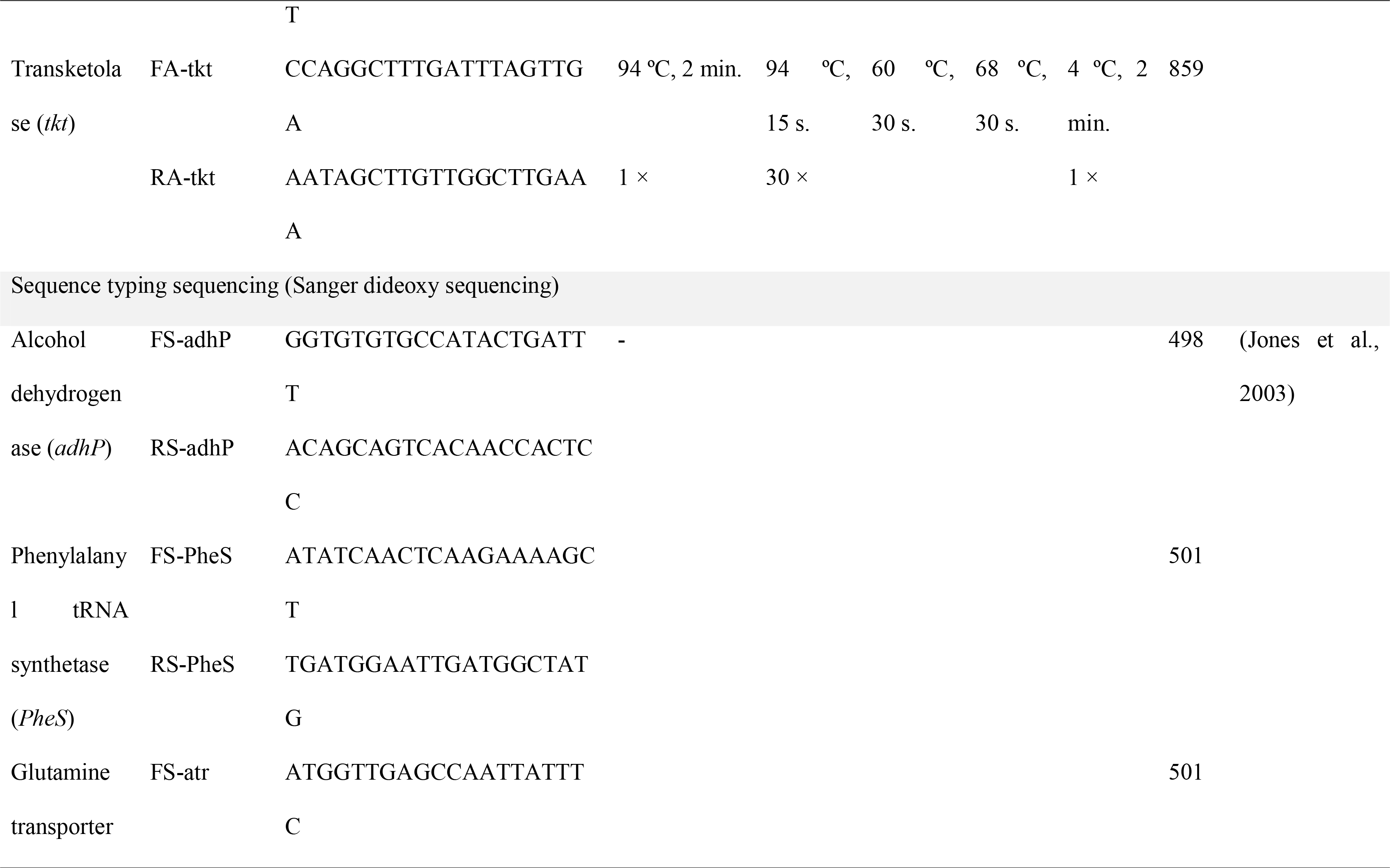
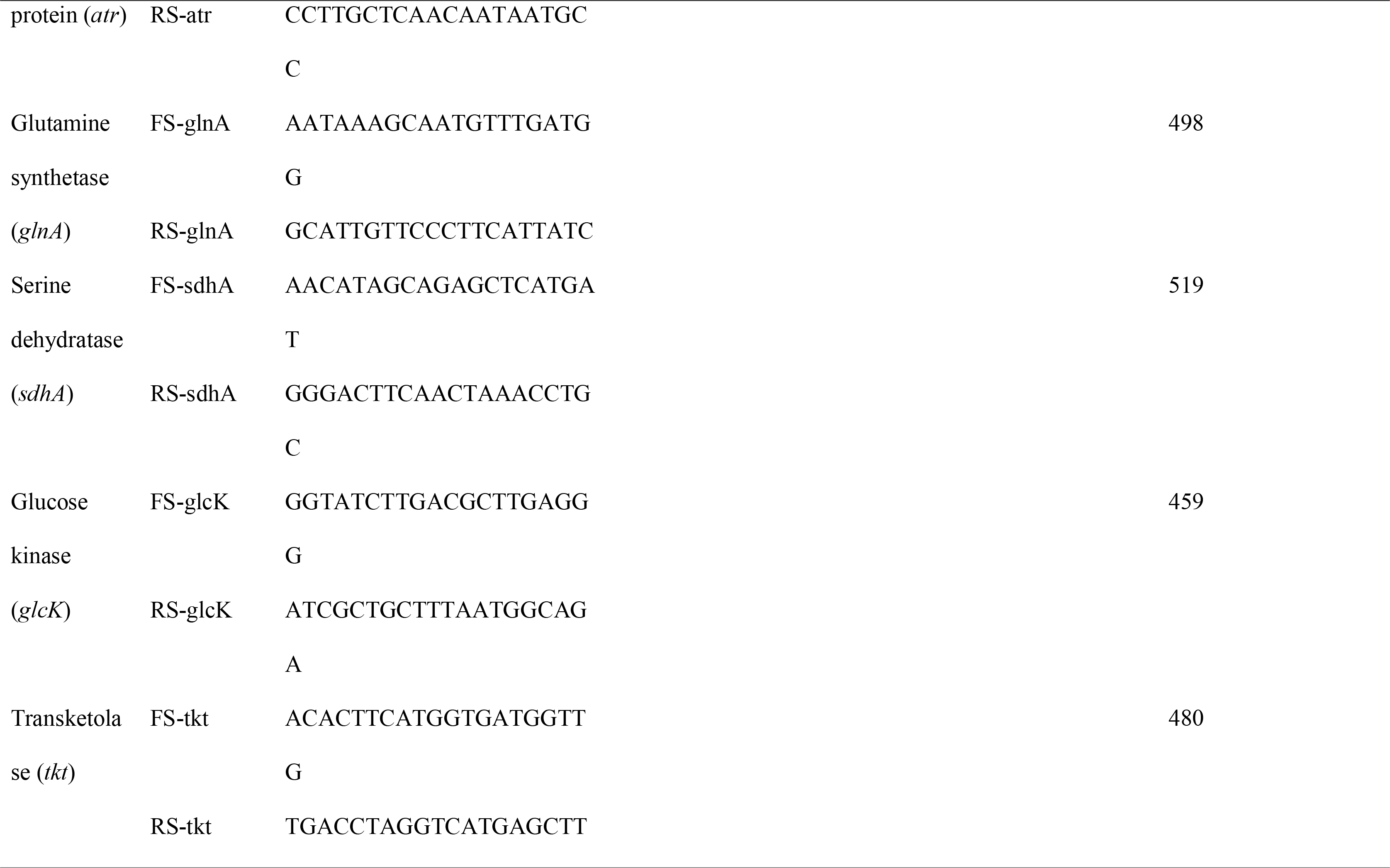
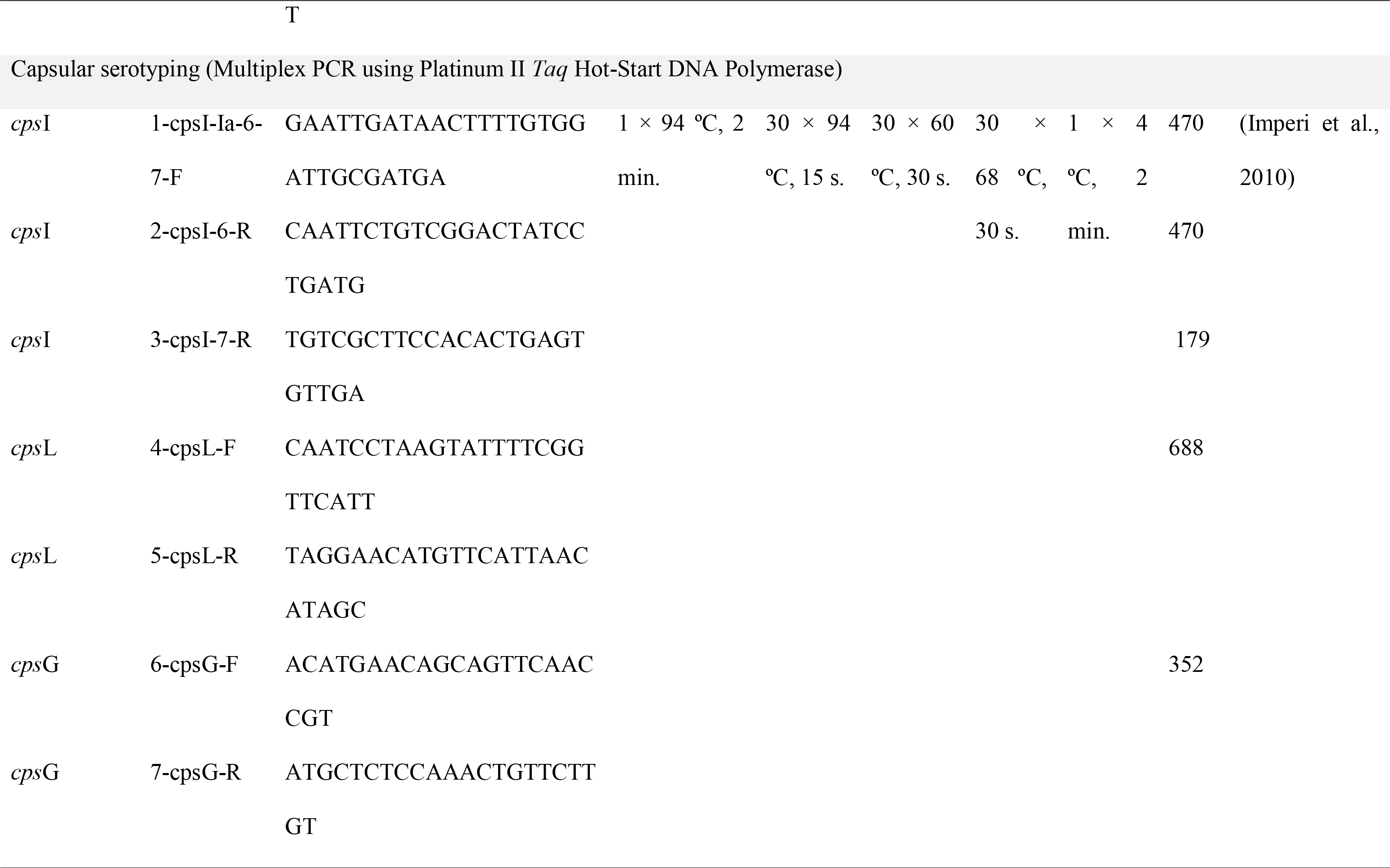
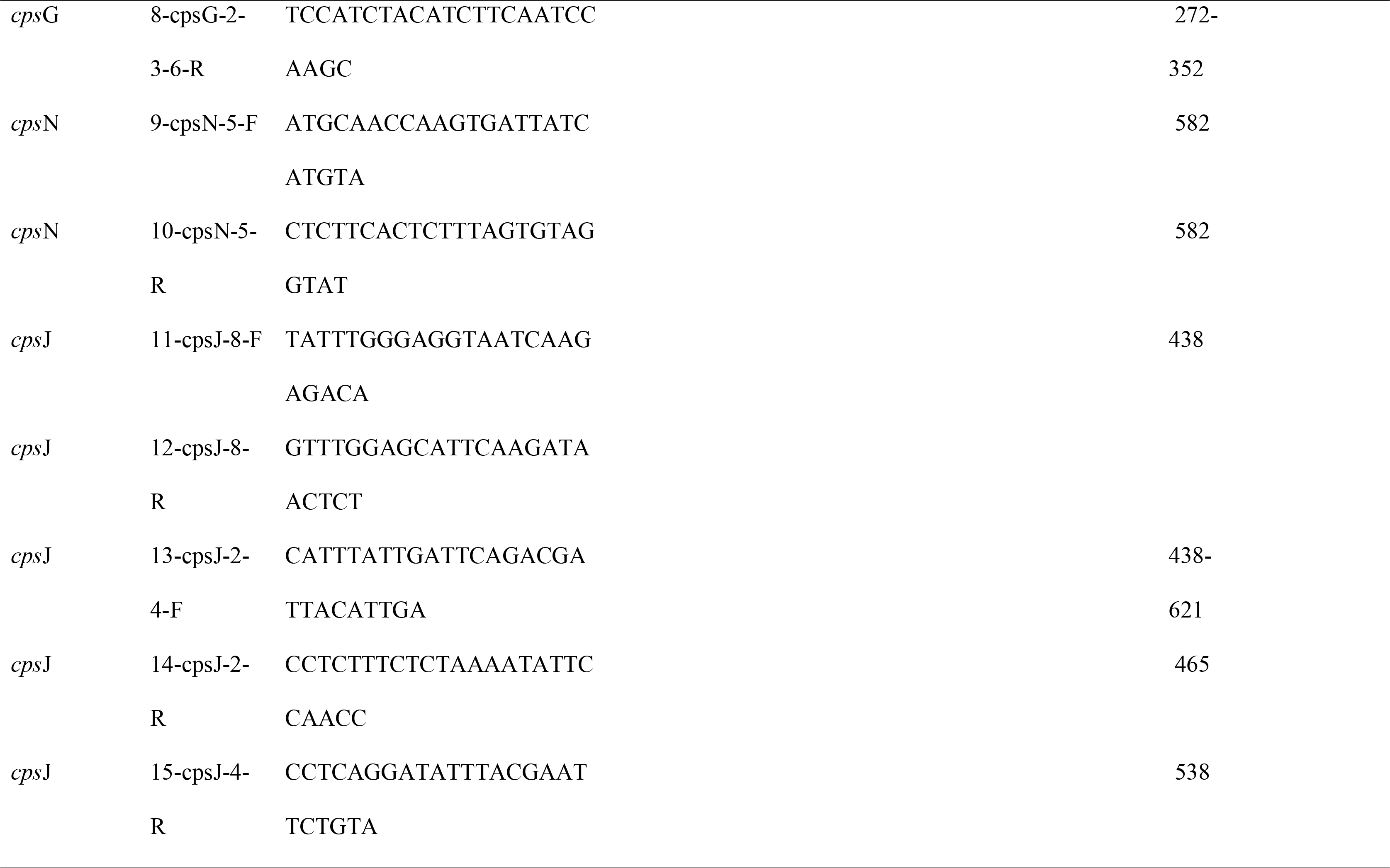
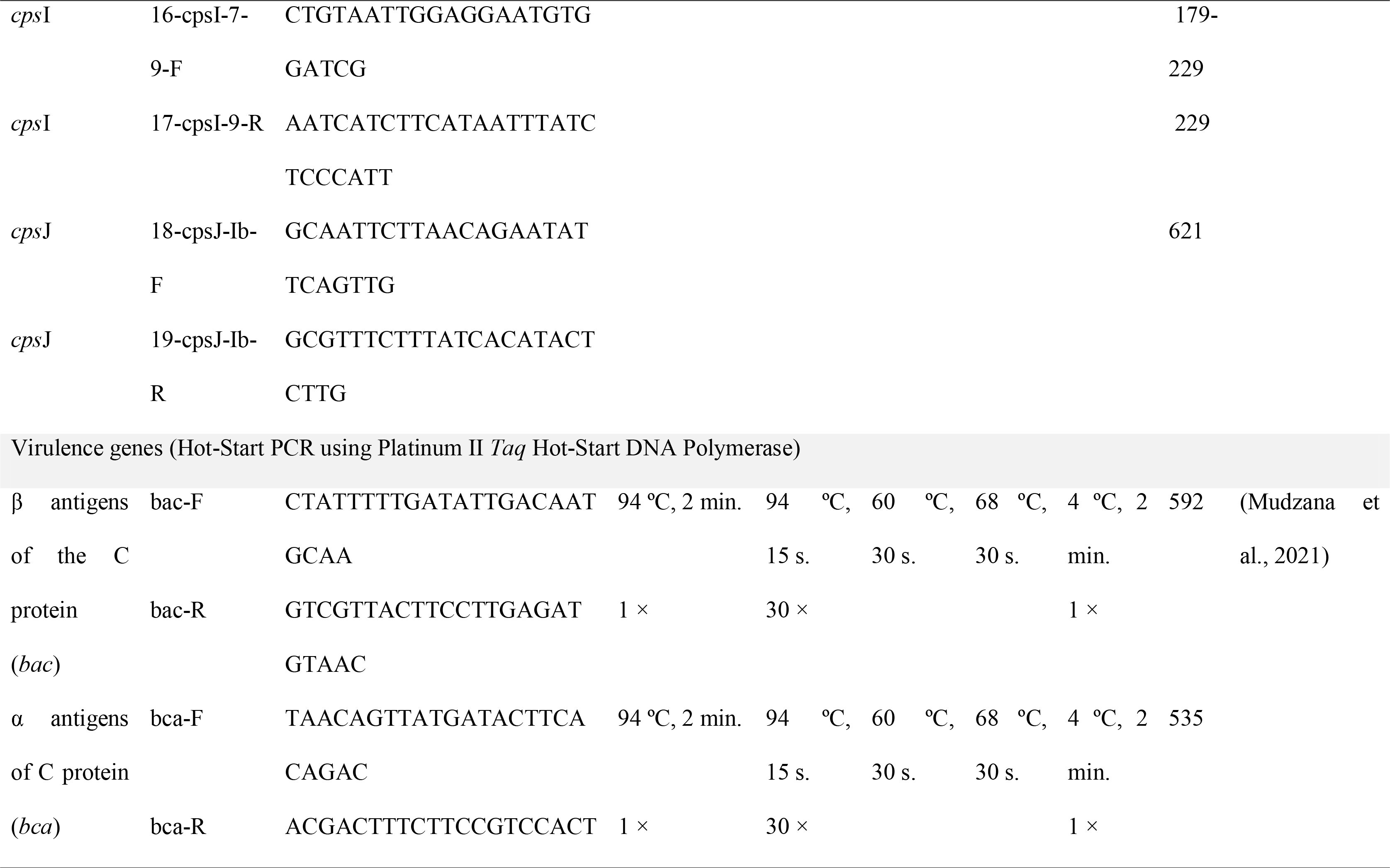
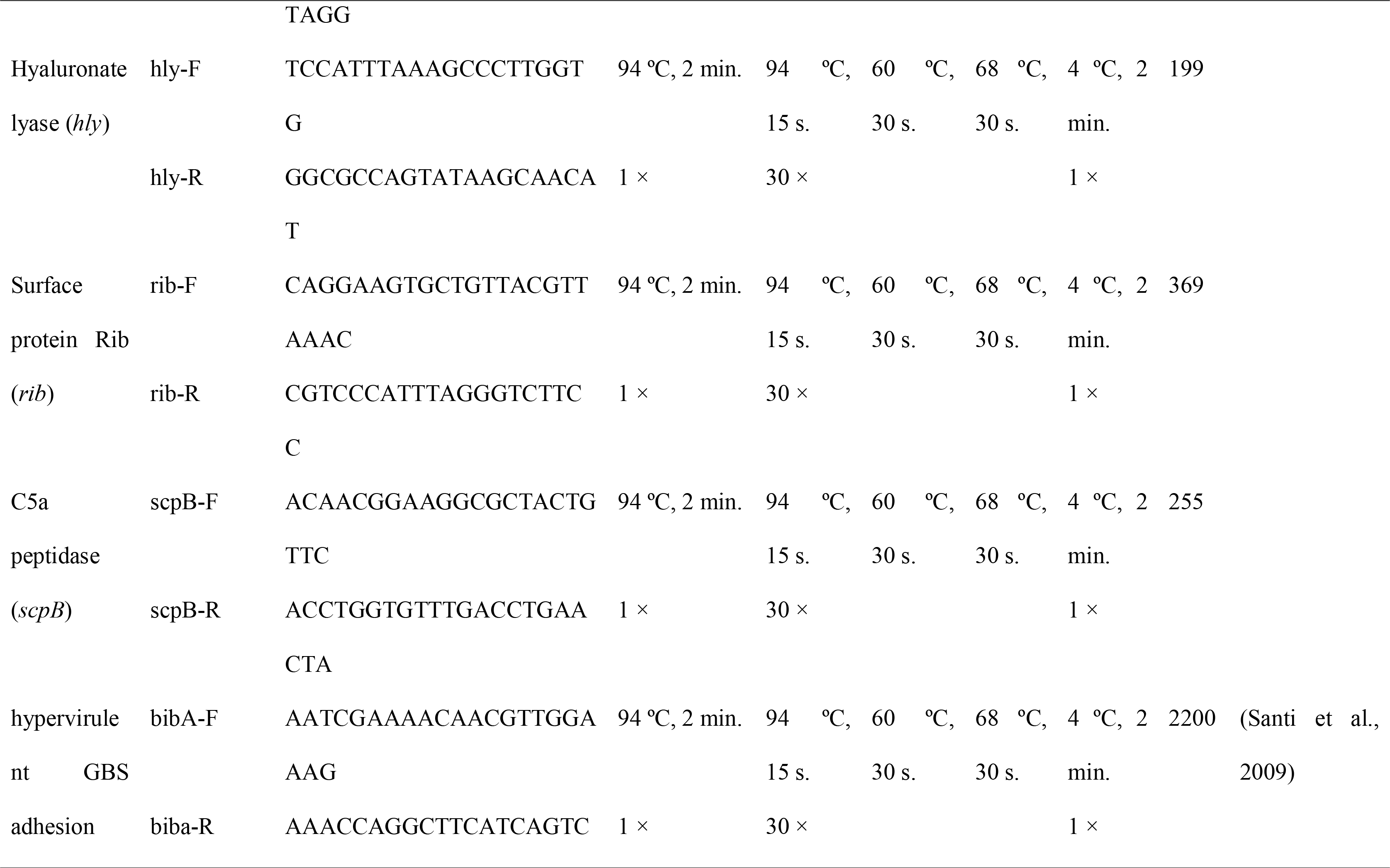
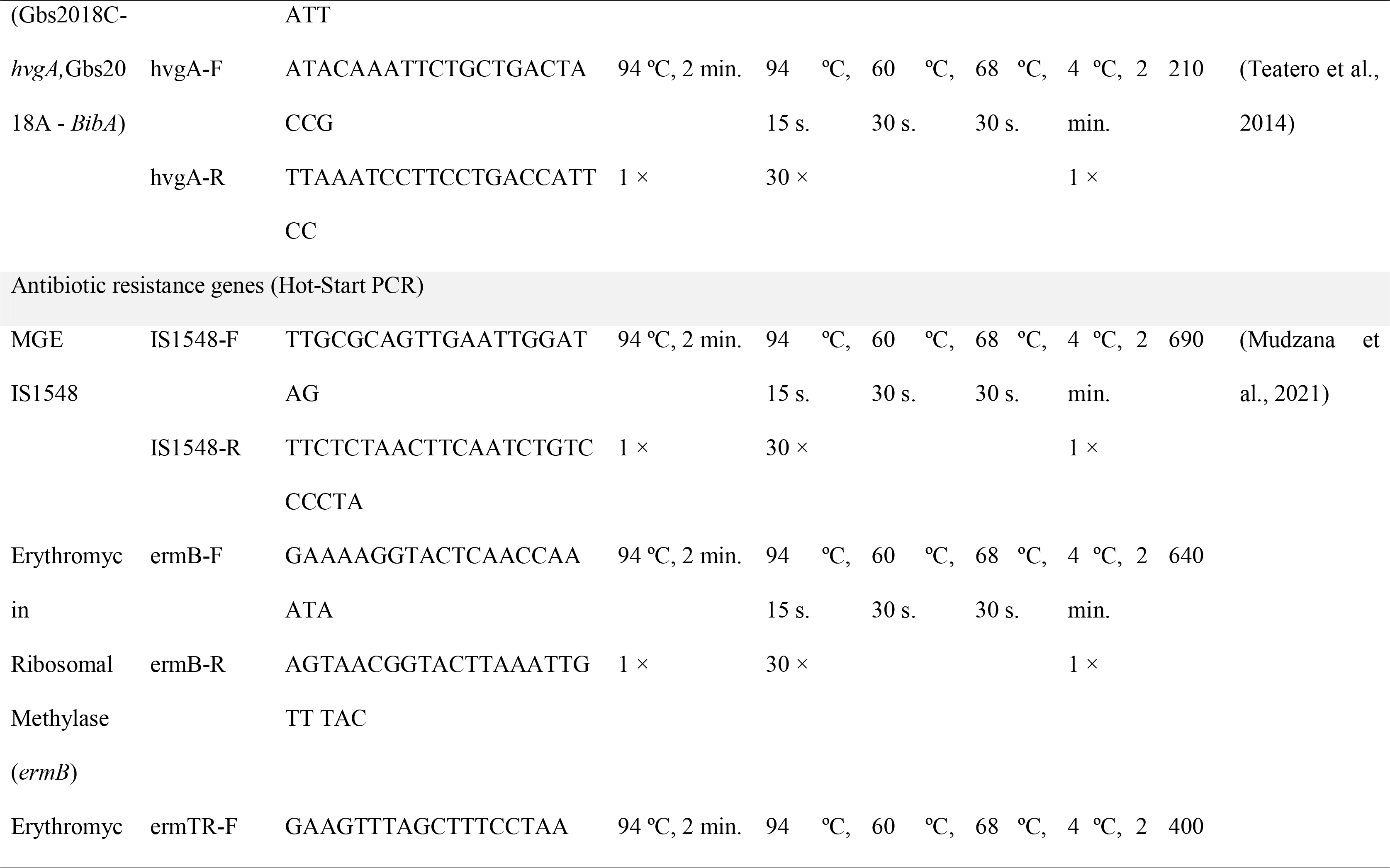
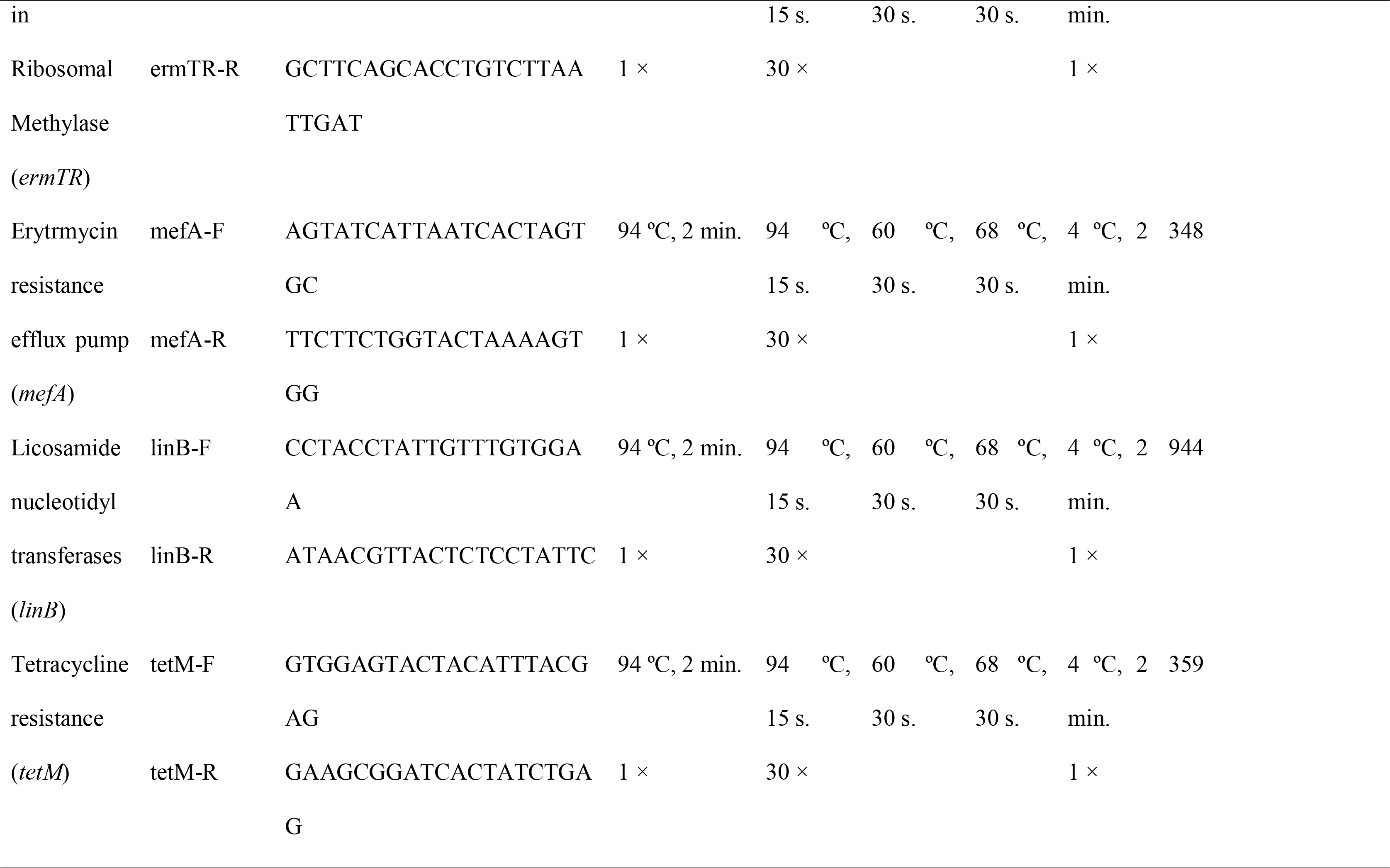
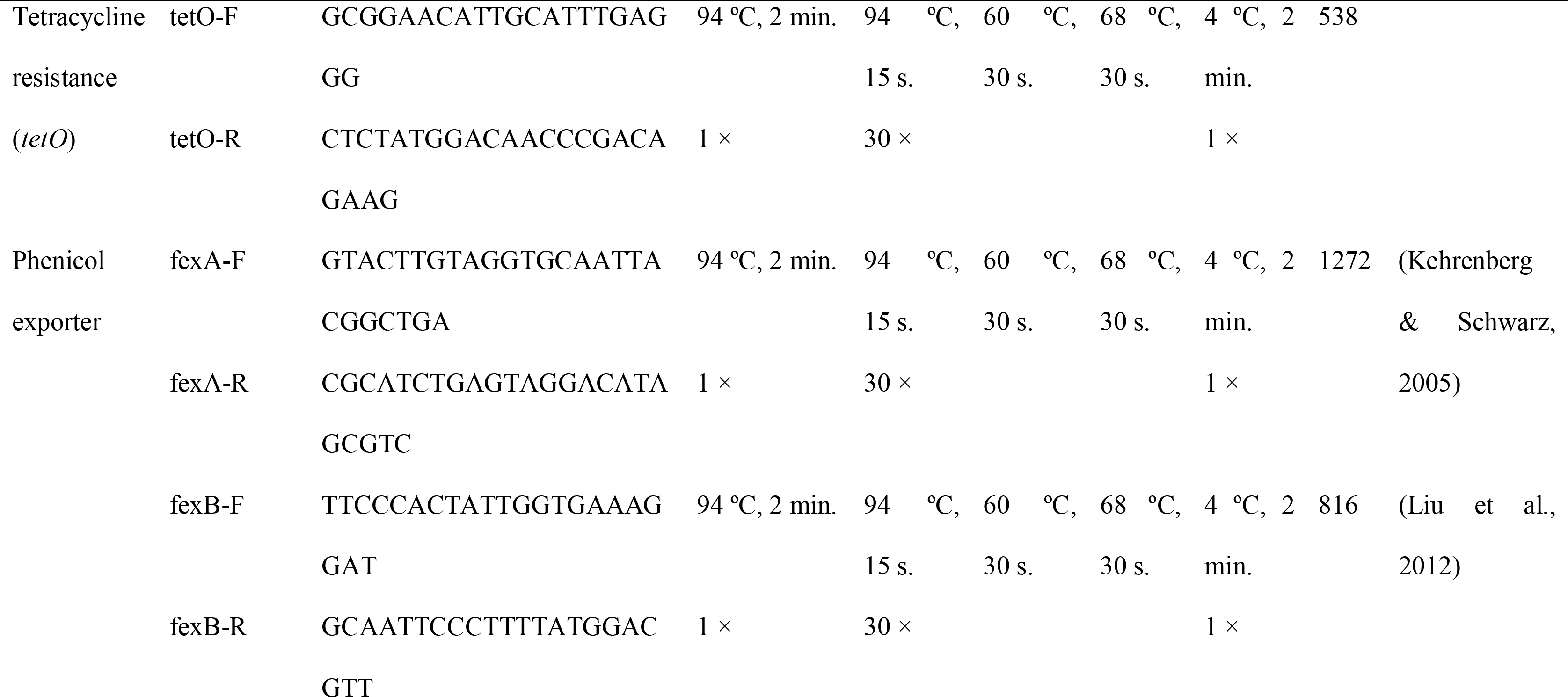
Primers and cycling conditions used for PCR amplification of different genes used for identification, multi-locus sequence typing (MLST), capsular serotyping, and detection of virulence and antibiotic resistance factors of 72 clinical isolates of Streptococcus agalactiae recovered between 2011 and 2018, from the brain, liver and kidney of diseased Nile tilapia, Oreochromis niloticus, farmed in different locations of Brazil.

#### 2.2.1. Sample preparation for PCR

Genomic DNA was extracted from the bacteria of freshly grown culture plates, using a DNeasy blood and tissue kit (Qiagen, Germany), following the manufacturer’s instructions (Qiagen, 2020). The DNA was quantified, using a NanoDrop™ One/One^C^ Microvolume UV- Vis Spectrophotometer (Thermo Fisher Scientific™, Brazil) and was used to perform Hot Start and Multiplex PCRs to amplify the *atr*, *cps*, *bac*, *bca*, *hly*, *rib*, *scpB*, IS1548, *ermB*, *ermTR*, *mefA*, *linB*, *tetM*, *tetO*, *fexA* and *fexB* genes, using the primers described in Table 1.

#### 2.2.2. Hot Start PCR

Hot Start PCR (25 µL reaction) was performed with 0.04 UµL^−1^ Invitrogen Platinum^TM^ II *Taq* Hot-Start DNA Polymerase (Thermo Fisher Scientific^TM^ Brazil, Lot 00813145), 1X Platinum^TM^ II PCR Buffer (Thermo Fisher Scientific^TM^ Brazil), 10 mM dNTP mix (Thermo Fisher Scientific^TM^ Brazil, Lot 1987090, 0.2 mM each), 0.2 µM forward primer, 0.2 µM reverse primer, extracted DNA (20–50 ng µ L^−1^ at a ratio of 260 280^−1^ between 1.8 and 2.0) and ultrapure water. Reactions (25 µL) were incubated in a thermal cycler (ProFlex PCR System, Thermo Fisher Scientific ^TM^ Brazil) under the conditions described in Table 1.

#### 2.2.3. Multiplex PCR

Multiplex PCR (25 µ L reaction) was performed as described in Table 1, with 0.3 U µ L^−1^ Invitrogen Platinum^TM^ II *Taq* Hot-Start DNA Polymerase (Thermo Fisher Scientific^TM^ Brazil, Lot 00813145), 1X Platinum^TM^ II PCR Buffer (Thermo Fisher Scientific^TM^ Brazil, Lot 01120395), 10 mM dNTP mix (Thermo Fisher Scientific^TM^ Brazil, Lot 1987090, 0.2 mM each), 0.25 µM of each primer except for primer 1 and 16 that were used at a concentration of 0.4 µM, extracted DNA (20–50 ng µ L^−1^ at a ratio of 260 280^−1^ between 1.8 and 2.0) and ultrapure water. Reactions (25 µ L) were incubated in a thermal cycler (ProFlex PCR System, Thermo Fisher Scientific ^TM^ Brazil) under the conditions described in Table 1.

#### 2.2.4. Electrophoresis

All PCR products were analysed by electrophoresis in a 1.5% agarose gel stained with 1µ L fluorescent stain for dsDNA (Nancy 520^®^, Sigma-Aldrich, Brazil), visualized under UV light, and sequenced as described in our previous study (Assane et al., 2021a).

### 2.3. Antimicrobial Susceptibility Testing (AST)

To determine the antimicrobial activity of florfenicol (FFC, ≥ 99.0% purity, Sigma-Aldrich), oxytetracycline (OTC, ≥ 95.0% purity, Supelco) and thiamphenicol (TAF, ≥ 99.0% purity, Sigma-Aldrich) against the clinical isolates, the broth microdilution method was performed as described in our previous studies (Assane et al., 2019, 2021b), following the Clinical Laboratory Standards Institute guideline VET03 (CLSI, 2020a) when possible. Briefly, antimicrobial stock solutions (1,024 µg mL^−1^ FFC, OTC or TAP) were prepared as described in our previous study (Assane et al., 2021b). The ranges of concentrations of antimicrobials used in the tests were 128–0.125 µg mL^−1^ FFC, OTC or TAP. *Streptococcus pneumoniae* ATCC^®^ 49619 and *Escherichia coli* ATCC 25922^®^ were used as quality controls.

Cation-adjusted Mueller-Hinton broth (CAMHB) supplemented with 4% (v/v) lysed horse blood (LHB) was used to perform the tests. For inoculum preparation, colony suspension (3–5 well-isolated colonies) of each strain tested was adjusted to 1–2 × 10^8^ CFU/mL using a spectrophotometer (0.08–0.13 at 625 nm) and diluted 1:150 (1 × 10^6^ CFU/mL). The diluted inoculum was immediately inoculated onto test plates containing the final twofold dilutions of each antimicrobial. The final inoculum concentration was 3.3–6.6 × 10^5^ CFU/mL, and the growth control only had the inoculum and CAMHB supplemented with 4% LHB.

Test plates were incubated in duplicate at 28°C ± 2°C for 44–48 h and minimum inhibitory concentration (MIC) endpoints were determined by visual inspection. Samples from three wells, the lowest concentration of antimicrobial without visible bacterial growth, were inoculated aseptically, in duplicate, onto CAMHA plates supplemented with sheep blood. Plates were incubated at 28°C ± 2°C for 44–48 hours to determine the MBC as the lowest concentration without bacterial growth.

The antimicrobial activity was categorized as bactericidal (MBC/MIC < 4) or bacteriostatic (MBC/MIC ≥ 4), based on the MBC-to-MIC ratio (Assane et al., 2019; Giguère et al., 2013). Due to the lack of standardized interpretative criteria for susceptibility testing of GBS isolated from aquatic organisms (CLSI, 2020a, 2020b), the strain categorization as wild-type or non-wild-type was avoided.

#### 2.3.1. Checkerboard Assay

To evaluate interactions between the studied antimicrobials against the clinical isolates, *in vitro* antimicrobial activity of their combinations was determined by checkerboard and time-kill kinetic analysis, using the MIC values determined previously (Leber, 2016). Test conditions (culture medium, incubation temperature and time, and inoculum concentration) were identical to those described in section. 2.3.. Antimicrobial concentration ranges on the test were determined based on the MIC of each antimicrobial against the tested strain.

MIC endpoints were determined by visual inspection and the fractional inhibitory concentration (FIC) was calculated as follows: FIC of antimicrobial A = MIC of antimicrobial A in combination/MIC of antimicrobial A alone, and FIC of antimicrobial B = MIC of antimicrobial B in combination/MIC of antimicrobial B alone. The FIC index (FICI) was defined as the sum of FICs. The interactions were classified as synergy (FICI ≤ 0.5), additive (0.5 < FICI ≤ 1), indifference (1 < FICI < 2) and antagonism (FICI ≥ 2) (EUCAST, 2000; Odds, 2003).

## 3. Results and discussion

Currently, *Streptococcus agalactiae*, group B *Streptococcus* (GBS), is the main bacterial pathogen of Nile tilapia, *Oreochromis niloticus*, and causes massive deaths in all stages of the farming cycle throughout the year, with higher frequency when the water temperature is higher (> 28 °C) (Asencios et al., 2016; Chideroli et al., 2017; Delannoy et al., 2021; Delphino et al., 2019; Verner-Jeffreys et al., 2018). Considering that isolates of GBS show distinct genetic diversity (Barony et al., 2017), as well as variability in biochemical and physiological characteristics (Goodfellow et al., 2012), information about their genetic diversity, virulence, presence of antibiotic resistance genes and antimicrobial susceptibility, are relevant for both epidemiological studies and the development of effective prevention and treatment strategies. In this study, we investigated the genetic diversity, virulence, presence of antibiotic resistance genes and antimicrobial susceptibility of 72 GBS linked to mass mortalities of cultured Nile tilapia in Brazil.

All clinical isolates studied were confirmed to be GBS, one (IA2022) from serotype III and 71 from serotype Ib, suggesting that serotype Ib was the most prevalent strain between 2011 and 2016 in the Paraná and São Paulo states (Brazil) (Table 2). Serotype III was only isolated from a disease outbreak in the Bahia state (Brazil) in 2018, corroborating the results from previous studies that indicate that this serotype mostly occurs in fish farms located in the northeastern region of Brazil (Leal, Queiroz, Pereira, Tavares, & Figueiredo, 2019). These serotypes, along with serotype Ia, are the most reported serotypes associated with disease outbreaks in different countries (Chen et al., 2015; Delannoy et al., 2021; Sudpraseart et al., 2021), including Brazil (Chideroli et al., 2017; Delphino et al., 2019; Godoy et al., 2013). Current data indicate that in Brazil, capsular serotype Ib and III, and ST-103, ST-260, ST-552, and ST-553 (CC552) are the most prevalent strains, although strains belonging to capsular serotype Ia and non-serotypeable strains have been isolated from disease outbreaks as well (Barony et al., 2017; Chideroli et al., 2017; Godoy et al., 2013).

**Table 2.**
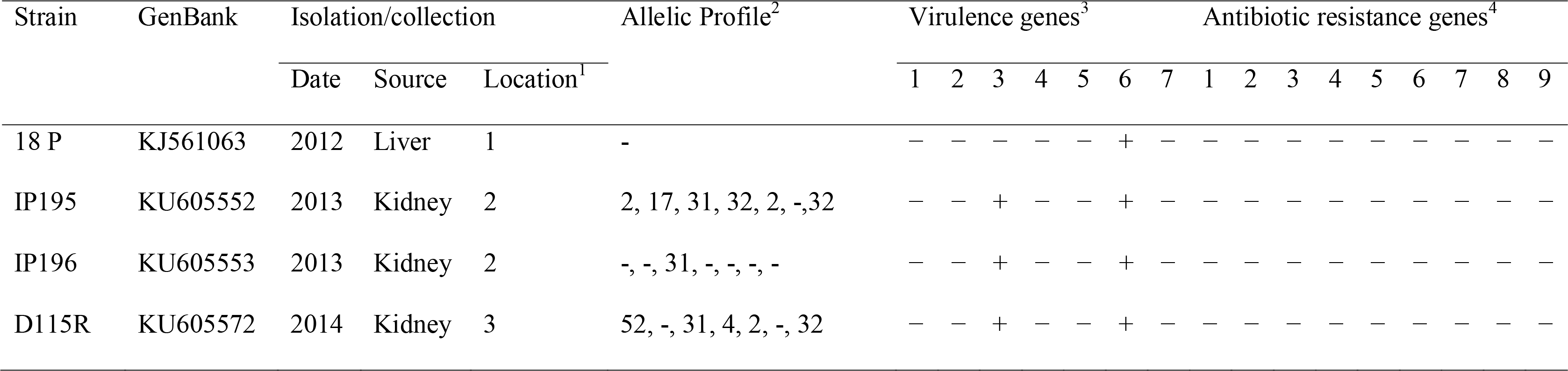
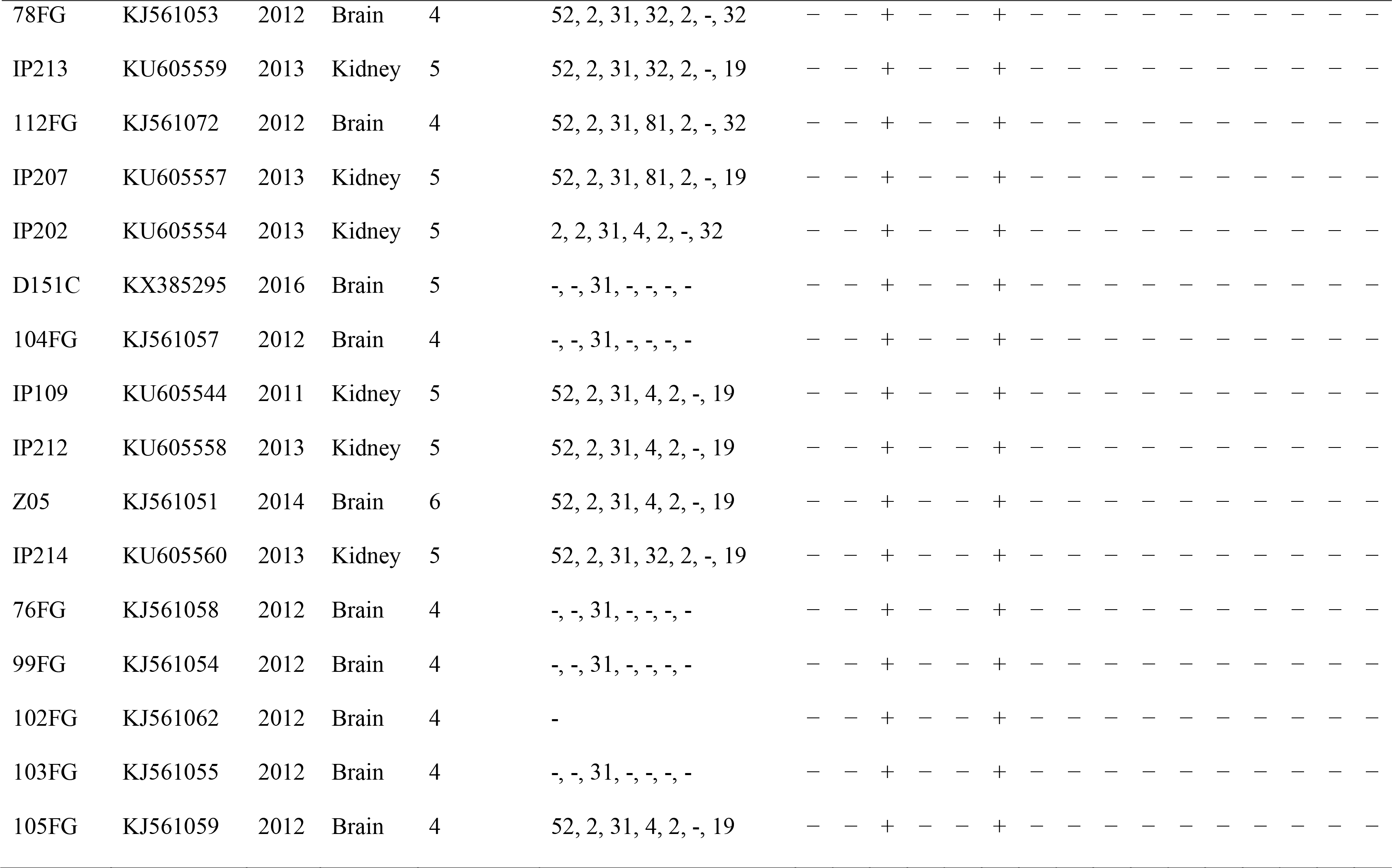
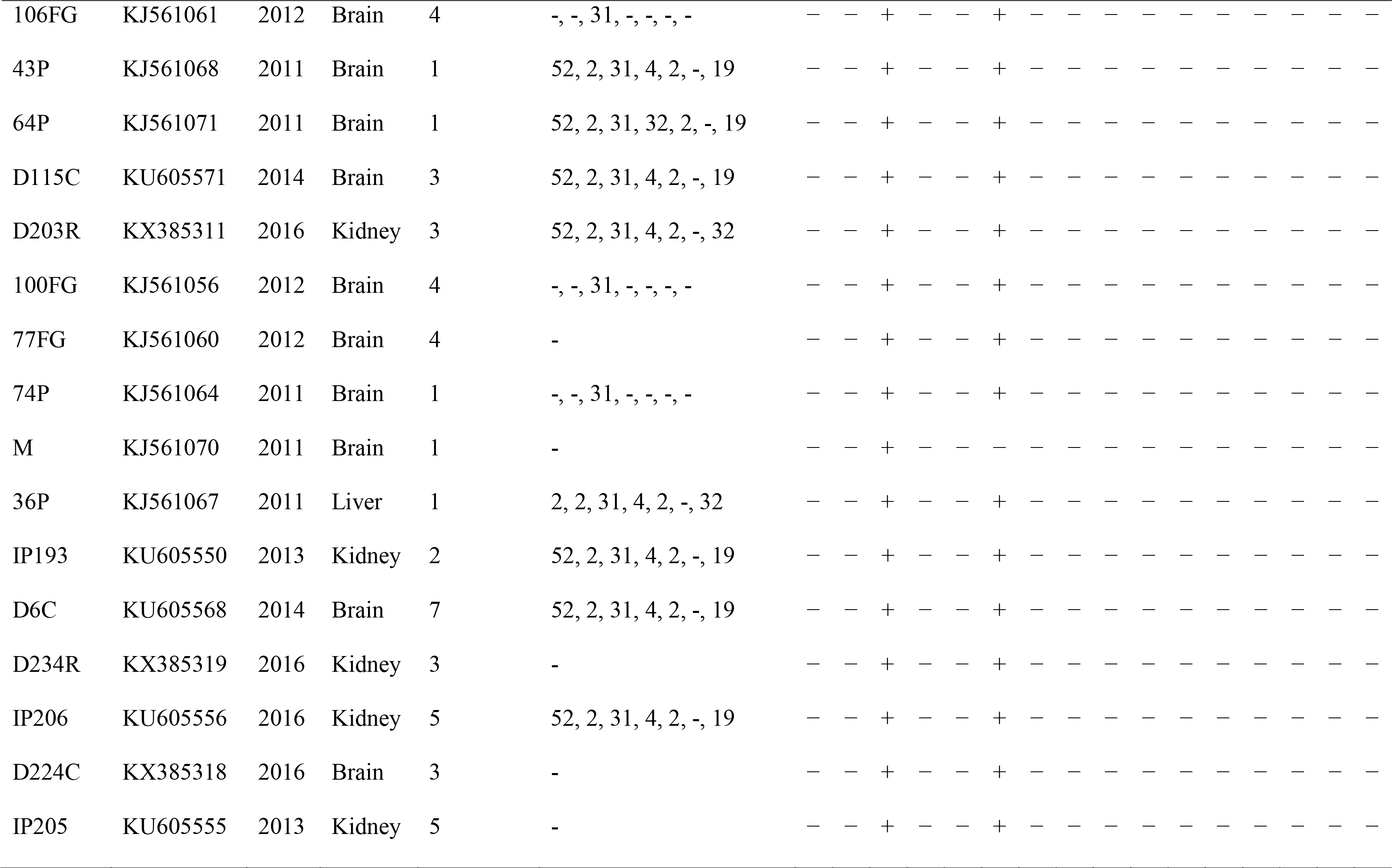
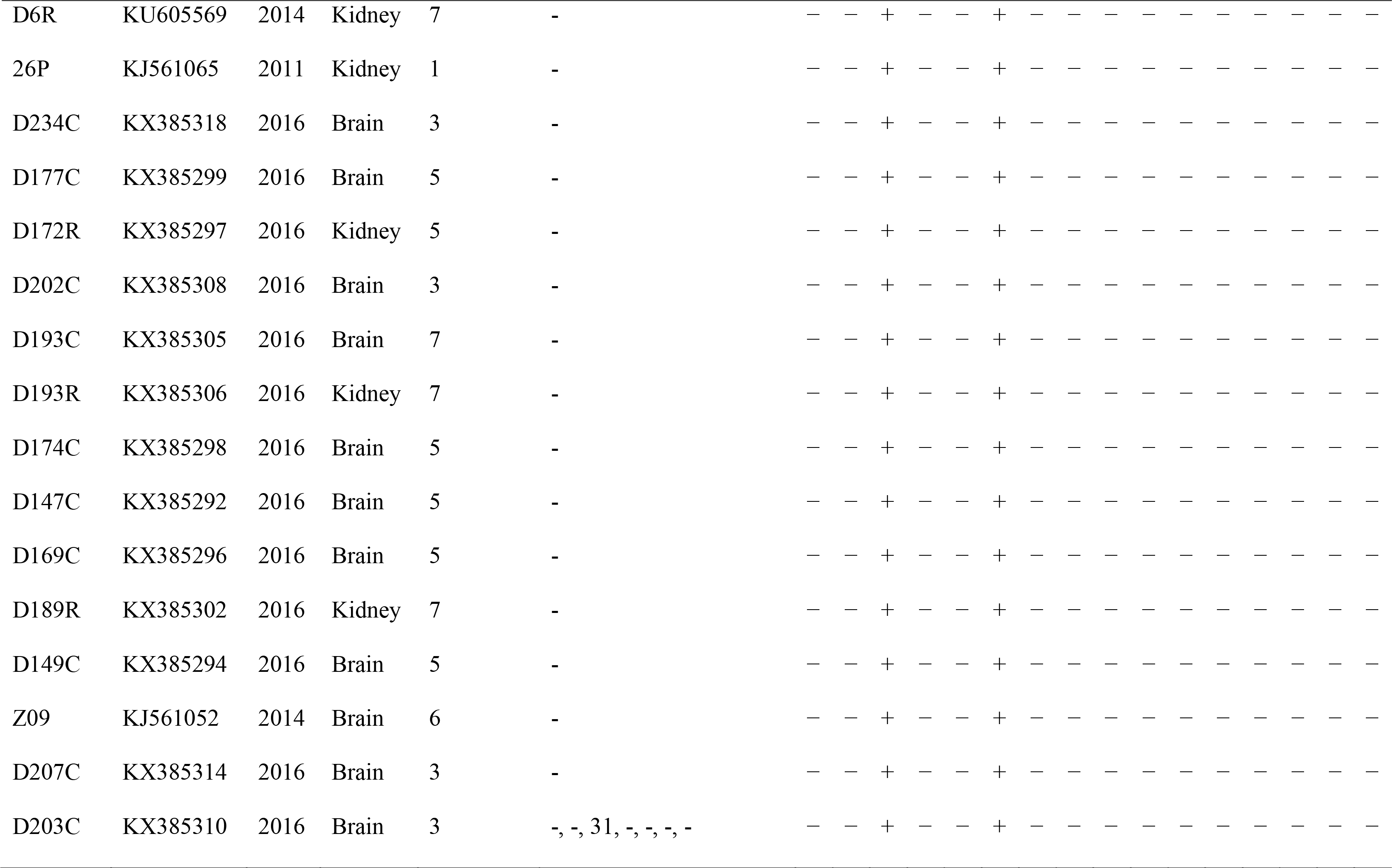
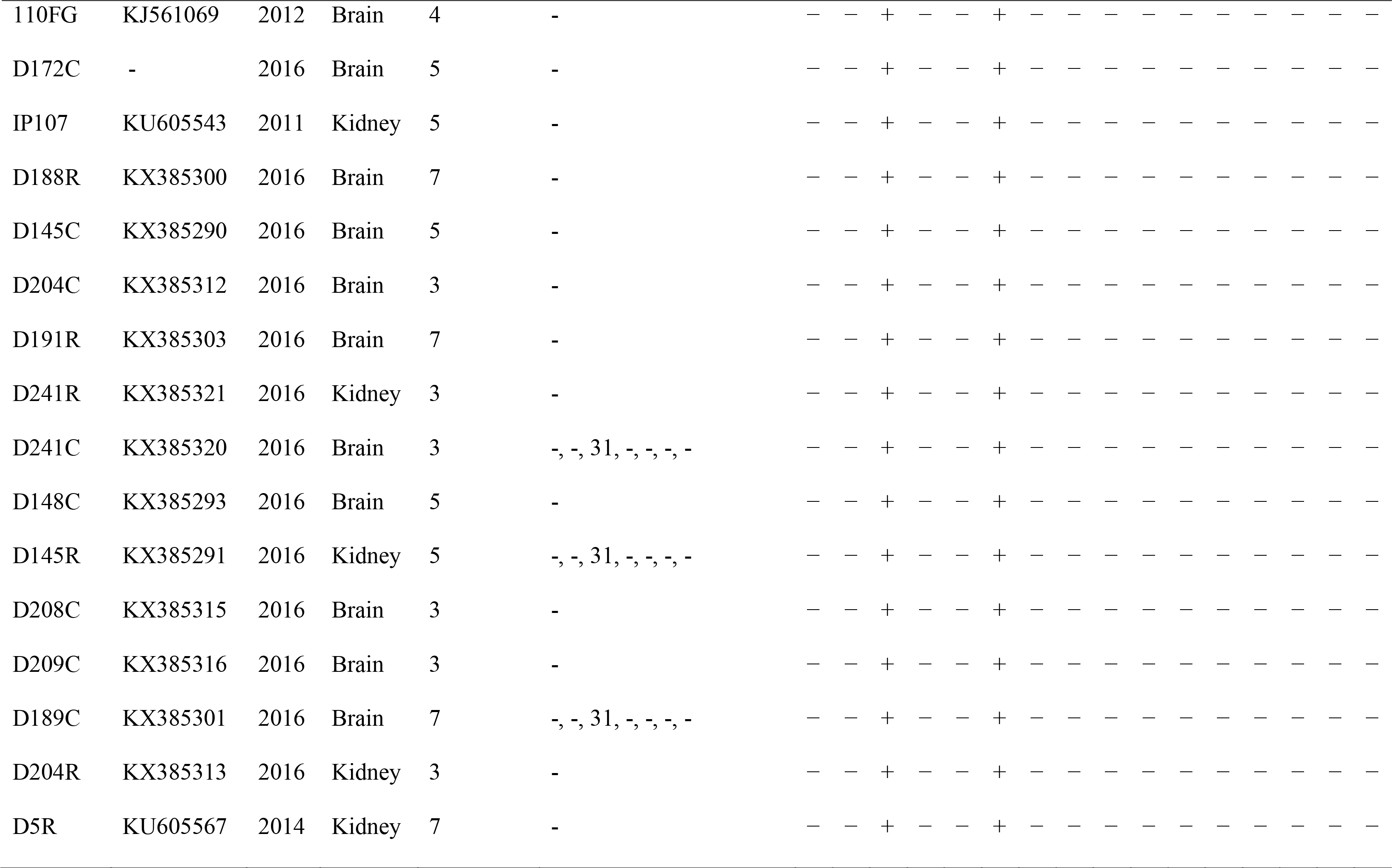
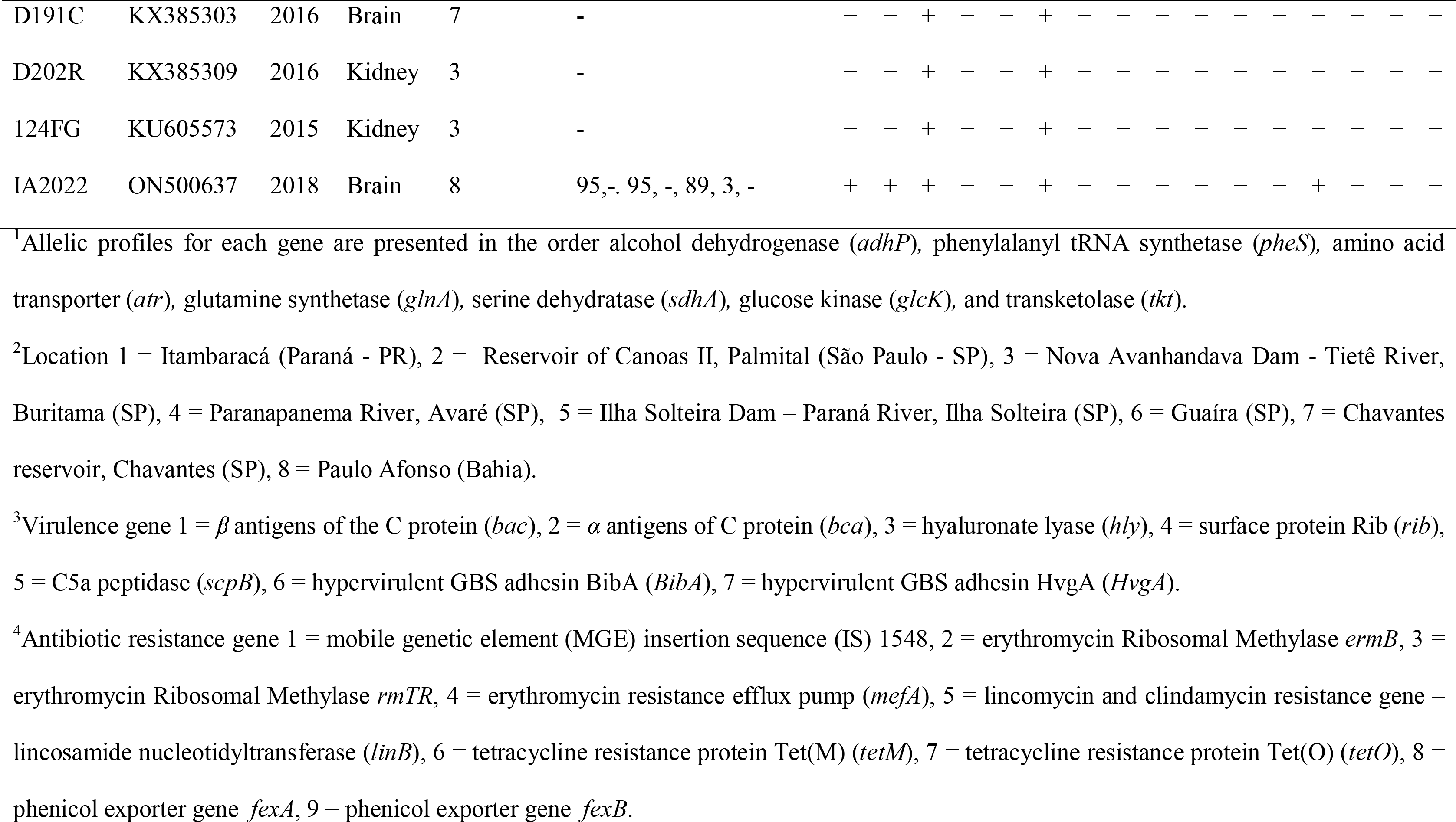
Allelic, virulence and antibiotic resistance factors profiles of 72 clinical isolates of Streptococcus agalactiae recovered between 2011 and 2018, from the internal organs of diseased Nile tilapia, Oreochromis niloticus, farmed in different locations of Brazil.

The allelic, virulence and antibiotic resistance factors profiles of each strain are presented in Table 2. The alleles defined for the MLST scheme were based on sequence lengths of between 242 and 652 bp for seven housekeeping genes (Table 3). Eight different allelic profiles (in the order: alcohol dehydrogenase - *adhP,* phenylalanyl tRNA synthetase - *pheS,* amino acid transporter - *atr,* glutamine synthetase - *glnA,* serine dehydratase - *sdhA,* glucose kinase - *glcK* and transketolase - *tkt*) were identified for the first time (2, 17, 31, 32, 2, -,32; 2, 2, 31, 4, 2, -, 32; 52, 2, 31, 32, 2, -, 19; 52, 2, 31, 32, 2, -, 32; 52, 2, 31, 4, 2, -, 19; 52, 2, 31, 4, 2, -, 32; 52, 2, 31, 81, 2, -, 19; and 52, 2, 31, 81, 2, -, 32), with 52, 2, 31, 4, 2, -, 19 being the most predominant. Due to the lack of reverse sequences of all alleles required for sequence type (ST) assignment (Jolley et al., 2018), these profiles were not deposited in the MLST database. Between one (*glcK*) and three (*adhP* and *glnA*) alleles were present at each locus. All strains, except IA2022, showed a partial gene deletion event on the *glcK* gene. This phenomenon was also reported in previous studies that evaluated the genetic diversity of *S. agalactiae*, which was isolated from fish cultured in Brazil (Assis, Tavares, Pereira, Figueiredo, & Leal, 2017), suggesting that most of our strains are genetically related to the strains previously reported in disease outbreaks in Brazil. This diversity of serotype Ib strains in the main tilapia production regions in Brazil, confirmed in the present study, may be the reason why commercial vaccines are ineffective in preventing streptococcosis, which is frequently reported by fish farmers.

**Table 3.**
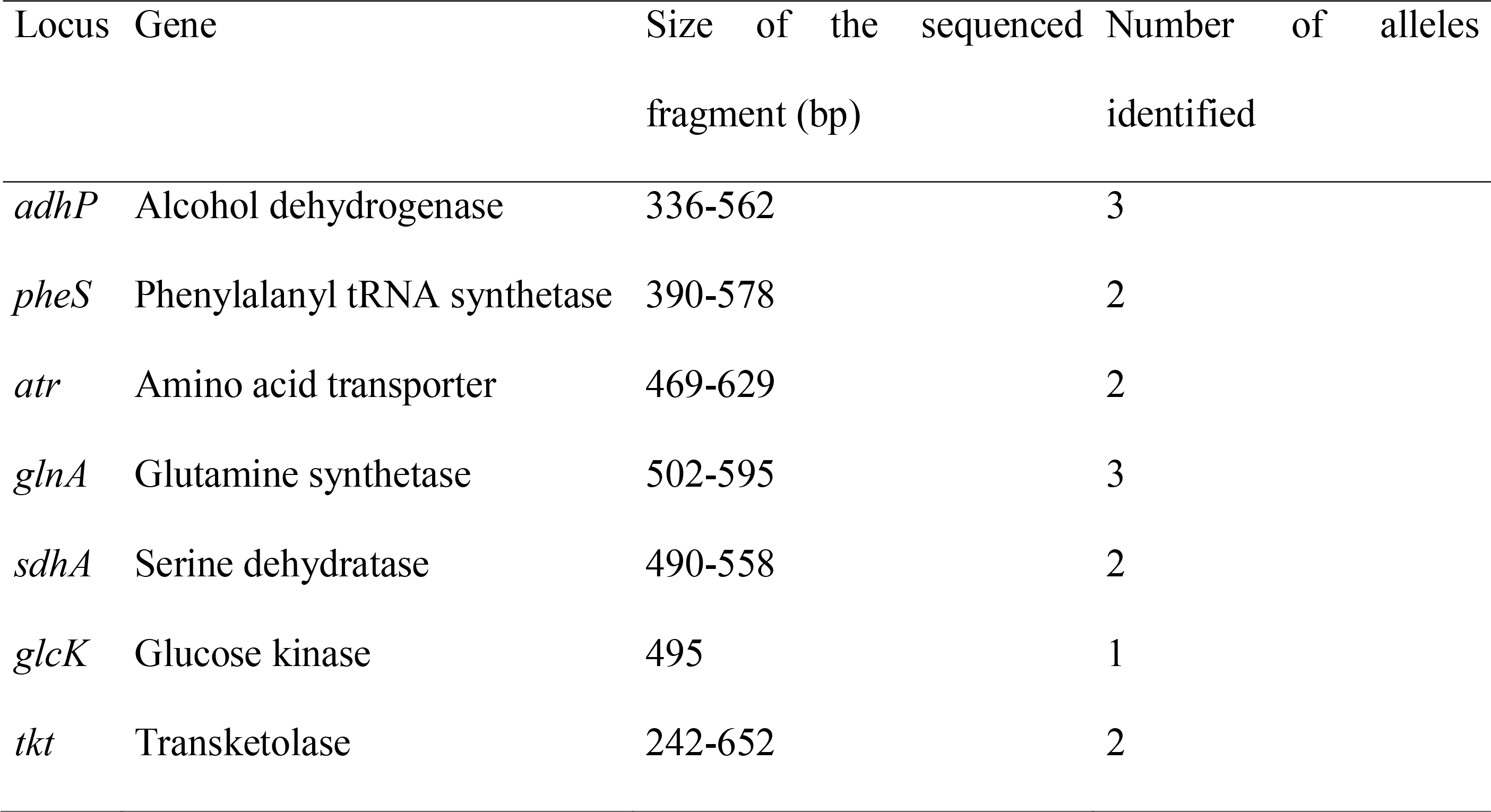
Characteristics of loci included in the GBS MLST system

| Locus | Gene | Size of the sequenced fragment (bp) | Number of alleles identified |
| --- | --- | --- | --- |
| *adhP* | Alcohol dehydrogenase | 336-562 | 3 |
| *pheS* | Phenylalanyl tRNA synthetase | 390-578 | 2 |
| *atr* | Amino acid transporter | 469-629 | 2 |
| *glnA* | Glutamine synthetase | 502-595 | 3 |
| *sdhA* | Serine dehydratase | 490-558 | 2 |
| *glcK* | Glucose kinase | 495 | 1 |
| *tkt* | Transketolase | 242-652 | 2 |

Surface protein Rib (*rib* = resistance to proteases, immunity, group B) and hypervirulent GBS adhesin BibA (*BibA*) were detected in all strains, except for 18P, which was negative for *rib* (Table 2). On the other hand, α and β antigens of the C protein (*bca* and *bac*, respectively) were only detected in IA2022. These results agree with previous studies which demonstrated that these proteins are associated with invasive strains that have been isolated from mammalians (Fluegge et al., 2011; Persson et al., 2008; Stålhammar-Carlemalm, Stenberg, & Lindahl, 1993) and may be evidence of an important role of these proteins in GBS infection in Nile tilapia. Considering that the expression of these antigens is correlated with protection (Bianchi-Jassir et al., 2020; Santi et al., 2009; Stålhammar-Carlemalm et al., 1993), our results indicate that all studied strains are good candidates to be used for the development of a protein and whole-cell vaccine against GBS.

Oxytetracycline (OTC) showed high minimum inhibitory concentration (MIC ≥ 16 µg/mL) values against several evaluated strains with negative results for tetracycline resistance genes (104FG, IP109, IP212, Z05, IP214, 43P, 64P, IP193, IP206, IP205, D174C, D147C, D149C, D207C, 110FG, D172C, IP107, D188R, D191R, D241R, D148C, D175R, D204R), indicating that the phenotypic antibiotic susceptibility patterns may not be similar to those obtained by genotyping. Similar results were observed with florfenicol (FFC) and thiamphenicol (TAP). However, the strain IA2022 was positive for the tetracycline resistance protein Tet(M) (*tetM*) and had a high MIC (32 µg/mL) of OTC (Table 4), corroborating the correlation between the resistance genotype determined by PCR assays and the antimicrobial susceptibility pattern. These results suggest that the higher MIC values of these strains, which were found to be negative by PCR, are probably associated with mechanisms not evaluated in this study and highlight the importance of antibiotic susceptibility testing for appropriate therapy decision-making.

**Table 4.** Antimicrobial activity (µg/mL) (minimum inhibitory concentration - MIC, minimum bactericidal concentration - MBC, and MBC/MIC ration) of florfenicol (FFC), thiamphenicol (TAP) and oxytetracycline (OTC) against 72 clinical isolates of Streptococcus agalactiae recovered between 2011 and 2018, from the internal organs of diseased Nile tilapia, Oreochromis niloticus, farmed in different locations of Brazil.

| Strain | GenBank | FFC |  | TAP |  | OTC |  | MBC/MIC |  |  | Interpretation* |  |  |
| --- | --- | --- | --- | --- | --- | --- | --- | --- | --- | --- | --- | --- | --- |
|  |  | MIC | MBC | MIC | MBC | MIC | MBC | FFC | TAP | OTC | FFC | TAP | OTC |
| 18P | KJ561063 | 2 | >8 | 16 | 16 | 0.5 | 2 | - | 1 | 4 | - | a | a |
| IP195 | KU605552 | 1 | 2 | 4 | 4 | 4 | 4 | 2 | 1 | 1 | a | a | a |
| IP196 | KU605553 | 2 | >8 | 4 | 16 | 4 | 4 | - | 4 | 1 | - | a | a |
| D115R | KU605572 | 2 | >8 | 16 | 16 | 0.5 | 0.5 | - | 1 | 1 | - | a | a |
| 78FG | KJ561053 | 0.25 | 0.25 | 16 | 16 | 0.25 | 0.25 | 1 | 1 | 1 | a | a | a |
| IP213 | KU605559 | 0.25 | 0.25 | 16 | >64 | 0.25 | 0.25 | 1 | - | 1 | a | - | a |
| 112FG | KJ561072 | 0.25 | 0.25 | 8 | 32 | 0.25 | 0.25 | 1 | 4 | 1 | a | a | a |
| IP207 | KU605557 | 1 | >4 | 4 | 4 | 0.25 | 0.25 | - | 1 | 1 | - | a | a |
| IP202 | KU605554 | 1 | >4 | 4 | 4 | 0.25 | 0.25 | - | 1 | 1 | - | a | a |
| D151C | KX385295 | <0.25 | - | 128 | 512 | 0.5 | 2 | - | 4 | 4 | - | a | a |
| 104FG | KJ561057 | 4 | 4 | 4 | 4 | >64 | - | 1 | 1 | - | a | a | - |
| IP109 | KU605544 | 2 | 2 | 8 | 32 | 256 | 256 | 1 | 4 | 1 | a | a | a |
| IP212 | KU605558 | 2 | 2 | 8 | 32 | 256 | 256 | 1 | 4 | 1 | a | a | a |
| Z05 | KJ561051 | 2 | 8 | 8 | 32 | 256 | 256 | 4 | 4 | 1 | a | a | a |
| IP214 | KU605560 | 2 | 8 | 8 | 32 | 128 | 128 | 4 | 4 | 1 | a | a | a |
| 76FG | KJ561058 | <0.25 | - | <0.25 | - | 0.5 | 0.5 | - | - | 1 | - | - | a |
| 99FG | KJ561054 | 8 | 8 | 32 | 32 | 0.5 | 0.5 | 1 | 1 | 1 | a | a | a |
| 102FG | KJ561062 | <0.25 | - | <0.25 | - | 0.5 | 0.5 | - | - | 1 | - | - | a |
| 103FG | KJ561055 | <0.25 | - | <0.25 | - | 0.5 | 0.5 | - | - | 1 | - | - | a |
| 105FG | KJ561059 | 4 | 4 | 4 | 4 | 2 | 2 | 1 | 1 | 1 | a | a | a |
| 106FG | KJ561061 | - | - | - | - | - | - | - | - | - | - | - | - |
| 43P | KJ561068 | 4 | 4 | 4 | 8 | 32 | 32 | 1 | 2 | 1 | a | a | a |
| 64P | KJ561071 | 2 | 8 | 64 | 64 | 16 | 16 | 4 | 1 | 1 | a | a | a |
| D115C | KU605571 | 4 | 8 | 128 | 512 | 4 | 16 | 2 | 4 | 2 | a | a | a |
| D203R | KX385311 | 2 | 8 | 8 | 32 | 2 | 2 | 4 | 4 | 1 | a | a | a |
| 100FG | KJ561056 | 4 | 8 | 4 | 4 | 1 | 1 | 2 | 1 | 1 | a | a | a |
| 77FG | KJ561060 | 4 | 8 | 4 | 4 | 8 | 8 | 2 | 1 | 1 | a | a | a |
| 74P | KJ561064 | 2 | 8 | 4 | 16 | 4 | 4 | 4 | 4 | 1 | a | a | a |
| M | KJ561070 | 64 | - | >256 | >256 | 8 | 8 | - | - | 1 | - | - | a |
| 36P | KJ561067 | 8 | 8 | 8 | 16 | 2 | 2 | 1 | 2 | 1 | a | a | a |
| IP193 | KU605550 | 4 | 8 | 32 | 128 | 64 | 64 | 2 | 4 | 1 | a | a | a |
| D6C | KU605568 | 2 | 2 | 32 | 128 | 1 | 1 | 1 | 4 | 1 | a | a | a |
| D234R | KX385319 | - | - | - | - | - | - | - | - | - | - | - | - |
| IP206 | KU605556 | 2 | 2 | 8 | 8 | 32 | 32 | 1 | 1 | 1 | a | a | a |
| D224C | KX385318 | - | - | - | - | - | - | - | - | - | - | - | - |
| IP205 | KU605555 | 2 | 4 | 4 | 4 | >256 | - | 2 | 1 | - | a | a | - |
| D6R | KU605569 | - | - | - | - | - | - | - | - | - | - | - | - |
| 26P | KJ561065 | - | - | - | - | - | - | - | - | - | - | - | - |
| D234C | KX385318 | 4 | 4 | 4 | 8 | 0.5 | 0.5 | 1 | 2 | 1 | a | a | a |
| D177C | KX385299 | 2 | 2 | 4 | 8 | 2 | 2 | 1 | 2 | 1 | a | a | a |
| D172R | KX385297 | 2 | 2 | 4 | 8 | 2 | 2 | 1 | 2 | 1 | a | a | a |
| D202C | KX385308 | 2 | 4 | 4 | 8 | 1 | 1 | 1 | 2 | 1 | a | a | a |
| D193C | KX385305 | 2 | 2 | 4 | 4 | 0.5 | 0.5 | 1 | 1 | 1 | a | a | a |
| D193R | KX385306 | 2 | 2 | 4 | 4 | 1 | 1 | 1 | 1 | 1 | a | a | a |
| D174C | KX385298 | 8 | 8 | 4 | 4 | >256 | - | 1 | 1 | - | a | a | - |
| D147C | KX385292 | 8 | 8 | 4 | 8 | >256 | - | - | - | - | - | - | - |
| D169C | KX385296 | 4 | 4 | 4 | 4 | 2 | 2 | - | 1 | 1 | - | a | a |
| D189R | KX385302 | 2 | 2 | 4 | 4 | 0.5 | 0.5 | - | 1 | 1 | - | a | a |
| D149C | KX385294 | 2 | 2 | 4 | 8 | >256 | - | - | 2 | - | - | a | - |
| Z09 | KJ561052 | 2 | 2 | 4 | 8 | 8 | 8 | - | 2 | 1 | - | a | a |
| D207C | KX385314 | 2 | 2 | 4 | 8 | 32 | 32 | - | 2 | 1 | - | a | a |
| D203C | KX385310 | 2 | 2 | 4 | 16 | 0.5 | 0.5 | - | 4 | 1 | - | a | a |
| 110FG | KJ561069 | 2 | 2 | 4 | 4 | 32 | 32 | - | 1 | 1 | - | a | a |
| D172C | - | 2 | 2 | 4 | 16 | 16 | 16 | - | 4 | 1 | - | a | a |
| IP107 | KU605543 | 2 | 2 | 4 | 4 | 16 | 16 | - | 1 | 1 | - | a | a |
| D188R | KX385300 | 2 | 2 | 4 | 4 | 16 | 16 | - | 1 | 1 | - | a | a |
| D145C | KX385290 | 2 | 2 | 4 | 16 | 2 | 2 | - | 4 | 1 | - | a | a |
| D204C | KX385312 | 2 | 2 | 4 | 8 | 1 | 1 | - | 2 | 1 | - | a | a |
| D191R | KX385303 | 2 | 8 | 4 | 16 | 16 | 16 | 4 | 4 | 1 | a | a | a |
| D241R | KX385321 | 2 | 4 | 4 | 16 | 32 | 32 | 2 | 4 | 1 | a | a | a |
| D241C | KX385320 | 2 | >8 | 4 | 16 | 8 | >32 | - | 4 | - | - | a | - |
| D148C | KX385293 | 2 | >8 | 4 | 16 | 32 | 32 | - | 4 | 1 | - | a | a |
| D145R | KX385291 | 2 | 8 | 4 | 16 | 32 | 32 | 4 | 4 | 1 | a | a | a |
| D208C | KX385315 | 2 | 8 | 4 | 8 | 1 | 2 | 4 | 2 | 2 | a | a | a |
| D209C | KX385316 | 2 | 8 | 4 | 16 | 4 | 8 | 4 | 4 | 2 | a | a | a |
| D189C | KX385301 | 2 | 4 | 4 | 4 | 0.5 | 1 | 2 | 1 | 2 | a | a | a |
| D204R | KX385313 | 2 | 4 | 4 | 16 | 32 | 32 | 2 | 4 | 1 | a | a | a |
| D5R | KU605567 | 2 | >8 | 4 | 16 | 4 | 4 | - | 4 | 1 | a | a | a |
| D191C | KX385303 | 2 | 8 | 4 | 8 | 1 | 4 | 4 | 2 | 4 | a | a | a |
| D202R | KX385309 | 2 | 4 | 4 | 4 | 0.5 | 0.5 | 2 | 1 | 1 | a | a | a |
| 124FG | KU605573 | 2 | 2 | 4 | 4 | 1 | >2 | 1 | 1 | - | a | a | - |
| IA2022 | ON500637 | 4 | 16 | 2 | 2 | 32 | 64 | 4 | 1 | 2 | a | a | a |

\* a = bactericidal, b = bacteriostatic.

All antimicrobials showed bactericidal activity against the tested strains, despite the high MIC values (≥ 16 µg/mL) observed against some strains (Table 4). Most strains showed higher MIC values (≥ 16 µg/mL) of OTC (23 strains) than FFC (one strain) and TAP (11 strains), indicating that FFC and TAP are more active than OTC against GBS.

The *in vitro* antimicrobial activity of TAP in combination with FFC or OTC, and FFC in combination with OTC, against clinical isolates with higher MIC values (≥ 16 µg/mL) against at least one of the combined antimicrobials is presented in Tables 5-7, respectively. The MIC of TAP against 64P (64 µg/mL) and D115C (128 µg/mL) were reduced fourfold and eightfold, respectively, after the TAP and FFC combination, resulting in synergic effects (ΣFIC = 0.375). However, the MIC values remained high (≥16 µg/mL), suggesting a low probability of therapeutic efficacy of the combination against these strains. On the other hand, the MIC of OTC against IA2022 (32 µg/mL) was reduced fourfold after the OTC combination with TAP or FFC, resulting in synergic effects (ΣFIC = 0.5) and a lower MIC value (8 µg/mL). Oxytetracycline combination with TAP or FFC also showed antagonistic activity against some strains (IP214, D148C and D241C). Despite indifference and antagonism being the most predominant activities in all combinations, the record of synergism, including in a strain with a resistance gene and phenotypic resistance, suggests that combination therapy can have therapeutic efficacy when well-planned. Previous studies also demonstrated the potential of this form of therapy against bacterial diseases in cultured fish (Assane, Gozi, Valladão, & Pilarski, 2019; Assane, Santos[Filho, et al., 2021; Bandeira Junior et al., 2018; Bandeira Junior et al., 2021; de Abreu Reis Ferreira et al., 2022). Our results suggest that the combination involving OTC and TAP or FFC is a likely candidate for improving the treatment of streptococcosis using combination therapy (CT), even for strains showing phenotypic and genotypic resistance to OTC. However, for the application of these combinations in aquaculture, further *in vivo* studies should be performed to determine their pharmacokinetics-pharmacodynamics properties and environmental and economic impacts on fish farms.

**Table 5.** *In vitro* antimicrobial activity of thiamphenicol (TAP) in combination with florfenicol (FFC) against clinical isolates of *Streptococcus agalactiae* recovered between 2011 and 2014, from the internal organs of diseased Nile tilapia, *Oreochromis niloticus*, farmed in different locations of Brazil.

| Strain | GenBank | MIC<sup>1</sup> alone (µg/mL) |  | MIC in combination (µg/mL) |  | Fold MIC in combination |  | ΣFIC<sup>2</sup> |
| --- | --- | --- | --- | --- | --- | --- | --- | --- |
|  |  | TAP | FFC | TAP | FFC | TAP | FFC |  |
| 18P | KJ561063 | 16.00 | 2.00 | 8.00 | 1.00 | 1/2 | 1/2 | 1.00 |
| D115R | KU605572 | 16.00 | 2.00 | 8.00 | 0.25 | 1/2 | 1/8 | 0.63 |
| 78FG | KJ561053 | 16.00 | 0.25 | 4.00 | 0.25 | 1/4 | 1 | 1.25 |
| IP213 | KU605559 | 16.00 | 0.25 | 8.00 | 0.25 | 1/2 | 1 | 1.50 |
| 99FG | KJ561054 | 32.00 | 8.00 | 8.00 | 4.00 | 1/4 | 1/2 | 0.75 |
| 64P | KJ561071 | 64.00 | 2.00 | 16.00 | 0.25 | 1/4 | 1/8 | 0.38 |
| D115C | KU605571 | 128.00 | 4.00 | 16.00 | 1.00 | 1/8 | 1/4 | 0.38 |
| IP193 | KU605550 | 32.00 | 4.00 | 16.00 | 2.00 | 1/2 | 1/2 | 1.00 |
| D6C | KU605568 | 32.00 | 2.00 | 8.00 | 2.00 | 1/4 | 1 | 1.25 |

<sup>1</sup>MIC = minimum inhibitory concentration.
<sup>2</sup>FIC = fractional inhibitory concentration.

**Table 6.** In vitro antimicrobial activity of thiamphenicol (TAP) in combination with oxytetracycline (OTC) against clinical isolates of Streptococcus agalactiae recovered between 2011 and 2018, from the internal organs of diseased Nile tilapia, Oreochromis niloticus, farmed in different locations of Brazil.

| Strain | GenBank | MIC<sup>1</sup> alone (ug/mL) |  | MIC in combination (µg/mL) |  | Fold MIC in combination |  | ΣFIC<sup>2</sup> |
| --- | --- | --- | --- | --- | --- | --- | --- | --- |
|  |  | TAP | OTC | TAP | OTC | TAP | OTC |  |
| 18P | KJ561063 | 16.00 | 0.50 | 8.00 | 0.50 | 1/2 | 1 | 1.50 |
| D115R | KU605572 | 16.00 | 0.50 | 8.00 | 0.50 | 1/2 | 1 | 1.50 |
| 78FG | KJ561053 | 16.00 | 0.25 | 4.00 | 0.13 | 1/4 | 1/2 | 0.75 |
| IP213 | KU605559 | 16.00 | 0.25 | 8.00 | 0.25 | 1/2 | 1 | 1.50 |
| D151C | KX385295 | 128.00 | 0.50 | 32.00 | 0.25 | 1/4 | 1/2 | 0.75 |
| IP109 | KU605544 | 8.00 | 256.00 | 4.00 | 128.00 | 1/2 | 1/2 | 1.00 |
| IP212 | KU605558 | 8.00 | 256.00 | 4.00 | 128.00 | 1/2 | 1/2 | 1.00 |
| Z05 | KJ561051 | 8.00 | 256.00 | 4.00 | 64.00 | 1/2 | 1/4 | 0.75 |
| IP214 | KU605560 | 8.00 | 128.00 | 4.00 | 64.00 | 1/2 | 1/2 | 1.00 |
| 99FG | KJ561054 | 32.00 | 0.50 | 8.00 | 0.25 | 1/4 | 1/2 | 0.75 |
| 43P | KJ561068 | 4.00 | 32.00 | 2.00 | 16.00 | 1/2 | 1/2 | 1.00 |
| 64P | KJ561071 | 64.00 | 16.00 | 16.00 | 8.00 | 1/4 | 1/2 | 0.75 |
| D115C | KU605571 | 128.00 | 4.00 | 16.00 | 2.00 | 1/8 | 1/2 | 0.63 |
| IP109 | KU605544 | 8.00 | 256.00 | 8.00 | 128.00 | 1 | 1/2 | 1.50 |
| IP212 | KU605558 | 8.00 | 256.00 | 8.00 | 128.00 | 1 | 1/2 | 1.50 |
| Z05 | KJ561051 | 8.00 | 256.00 | 8.00 | 128.00 | 1 | 1/2 | 1.50 |
| IP214 | KU605560 | 8.00 | 128.00 | 8.00 | 128.00 | 1 | 1 | 2.00 |
| IP193 | KU605550 | 32.00 | 64.00 | 16.00 | 16.00 | 1/2 | 1/4 | 0.75 |
| D6C | KU605568 | 32.00 | 1.00 | 8.00 | 0.50 | 1/4 | 1/2 | 0.75 |
| IP206 | KU605556 | 8.00 | 32.00 | 8.00 | 8.00 | 1 | 1/4 | 1.25 |
| D207C | KX385314 | 4.00 | 32.00 | 2.00 | 16.00 | 1/2 | 1/2 | 1.00 |
| 110FG | KJ561069 | 4.00 | 32.00 | 2.00 | 8.00 | 1/2 | 1/4 | 0.75 |
| D172C | - | 4.00 | 16.00 | 2.00 | 8.00 | 1/2 | 1/2 | 1.00 |
| D191R | KX385303 | 4.00 | 16.00 | 2.00 | 8.00 | 1/2 | 1/2 | 1.00 |
| D241R | KX385321 | 4.00 | 32.00 | 2.00 | 32.00 | 1/2 | 1 | 1.50 |
| D148C | KX385293 | 4.00 | 32.00 | 4.00 | 32.00 | 1 | 1 | 2.00 |
| D145R | KX385291 | 4.00 | 32.00 | 2.00 | 16.00 | 1/2 | 1/2 | 1.00 |
| D204R | KX385313 | 4.00 | 32.00 | 2.00 | 16.00 | 1/2 | 1/2 | 1.00 |
| IA2022 | ON500637 | 2.00 | 32.00 | 0.50 | 8.00 | 1/4 | 1/4 | 0.50 |

<sup>1</sup>MIC = minimum inhibitory concentration.
<sup>2</sup>FIC = fractional inhibitory concentration.

**Table 7.** *In vitro* antimicrobial activity of florfenicol (FFC) in combination with oxytetracycline (OTC) against clinical isolates of *Streptococcus agalactiae* recovered between 2011 and 2018, from the internal organs of diseased Nile tilapia, *Oreochromis niloticus*, farmed in different locations of Brazil.

| Strain | GenBank | MIC alone (µg/mL) |  | MIC in combination (µg/mL) |  | Fold MIC in combination |  | ΣFIC |
| --- | --- | --- | --- | --- | --- | --- | --- | --- |
|  |  | FFC | OTC | FFC | OTC | FFC | OTC |  |
| IP109 | KU605544 | 2.00 | 256.00 | 1.00 | 128.00 | 1/2 | 1/2 | 1.00 |
| IP212 | KU605558 | 2.00 | 256.00 | 0.25 | 128.00 | 1/8 | 1/2 | 0.63 |
| Z05 | KJ561051 | 2.00 | 256.00 | 1.00 | 128.00 | 1/2 | 1/2 | 1.00 |
| IP214 | KU605560 | 2.00 | 128.00 | 0.25 | 64.00 | 1/8 | 1/2 | 0.63 |
| 43P | KJ561068 | 4.00 | 32.00 | 2.00 | 16.00 | 1/2 | 1/2 | 1.00 |
| 64P | KJ561071 | 2.00 | 16.00 | 0.25 | 8.00 | 1/8 | 1/2 | 0.63 |
| IP109 | KU605544 | 2.00 | 256.00 | 1.00 | 128.00 | 1/2 | 1/2 | 1.00 |
| IP212 | KU605558 | 2.00 | 256.00 | 1.00 | 128.00 | 1/2 | 1/2 | 1.00 |
| Z05 | KJ561051 | 2.00 | 256.00 | 1.00 | 64.00 | 1/2 | 1/4 | 0.75 |
| IP214 | KU605560 | 2.00 | 128.00 | 1.00 | 16.00 | 1/2 | 1/8 | 0.63 |
| IP193 | KU605550 | 4.00 | 64.00 | 2.00 | 16.00 | 1/2 | 1/4 | 0.75 |
| IP206 | KU605556 | 2.00 | 32.00 | 1.00 | 8.00 | 1/2 | 1/4 | 0.75 |
| D207C | KX385314 | 2.00 | 32.00 | 1.00 | 16.00 | 1/2 | 1/2 | 1.00 |
| 110FG | KJ561069 | 2.00 | 32.00 | 1.00 | 8.00 | 1/2 | 1/4 | 0.75 |
| D172C | - | 4.00 | 16.00 | 4.00 | 8.00 | 1 | 1/2 | 1.50 |
| D191R | KX385303 | 2.00 | 16.00 | 2.00 | 8.00 | 1 | 1/2 | 1.50 |
| D241R | KX385321 | 2.00 | 32.00 | 2.00 | 32.00 | 1 | 1 | 2.00 |
| D148C | KX385293 | 2.00 | 32.00 | 2.00 | 32.00 | 1 | 1 | 2.00 |
| D145R | KX385291 | 2.00 | 32.00 | 2.00 | 16.00 | 1 | 1/2 | 1.50 |
| D204R | KX385313 | 2.00 | 32.00 | 2.00 | 16.00 | 1 | 1/2 | 1.50 |
| IA2022 | ON500637 | 4.00 | 32.00 | 1.00 | 8.00 | 1/4 | 1/4 | 0.50 |

<sup>1</sup>MIC = minimum inhibitory concentration, <sup>2</sup>FIC = fractional inhibitory concentration.

In conclusion, our findings revealed that genetically diverse virulent *S. agalactiae* serotype Ib is the most common and widely distributed in Nile tilapia, which is cultured in the southern and southeastern regions of Brazil, and due to its economic impacts and high MIC values of the available antimicrobials against it, the authorities and aquaculture workers should be more concerned about the prudent use of antimicrobials in fish farms and find eco- friendly and more effective preventive and therapeutic options. Therefore, further studies should be performed to develop effective prophylactic and therapeutic strategies against GBS.

## Acknowledgements

The authors thank the São Paulo Research Foundation (FAPESP) (grant 2019/22775-0 and 2019/23029-0) and Coordenação de Aperfeiçoamento de Pessoal de Nível Superior – Brasil (CAPES) [Finance code 001].

